# Determinants of renin cell differentiation: a single cell epi-transcriptomics approach

**DOI:** 10.1101/2023.01.18.524595

**Authors:** Alexandre G Martini, Jason P. Smith, Silvia Medrano, Nathan C. Sheffield, Maria Luisa S. Sequeira-Lopez, R. Ariel Gomez

## Abstract

**Rationale:** Renin cells are essential for survival. They control the morphogenesis of the kidney arterioles, and the composition and volume of our extracellular fluid, arterial blood pressure, tissue perfusion, and oxygen delivery. It is known that renin cells and associated arteriolar cells descend from *FoxD1*+ progenitor cells, yet renin cells remain challenging to study due in no small part to their rarity within the kidney. As such, the molecular mechanisms underlying the differentiation and maintenance of these cells remain insufficiently understood.

**Objective:** We sought to comprehensively evaluate the chromatin states and transcription factors (TFs) that drive the differentiation of *FoxD1*+ progenitor cells into those that compose the kidney vasculature with a focus on renin cells.

**Methods and Results:** We isolated single nuclei of *FoxD1*+ progenitor cells and their descendants from *FoxD1*^*cre/+*^;*R26R-mTmG* mice at embryonic day 12 (E12) (n_cells_=1234), embryonic day 18 (E18) (n_cells_=3696), postnatal day 5 (P5) (n_cells_=1986), and postnatal day 30 (P30) (n_cells_=1196). Using integrated scRNA-seq and scATAC-seq we established the developmental trajectory that leads to the mosaic of cells that compose the kidney arterioles, and specifically identified the factors that determine the elusive, myo-endocrine adult renin-secreting juxtaglomerular (JG) cell. We confirm the role of *Nfix* in JG cell development and renin expression, and identified the myocyte enhancer factor-2 (MEF2) family of TFs as putative drivers of JG cell differentiation.

**Conclusions:** We provide the first developmental trajectory of renin cell differentiation as they become JG cells in a single-cell atlas of kidney vascular open chromatin and highlighted novel factors important for their stage-specific differentiation. This improved understanding of the regulatory landscape of renin expressing JG cells is necessary to better learn the control and function of this rare cell population as overactivation or aberrant activity of the RAS is a key factor in cardiovascular and kidney pathologies.

## Introduction

Renin cells emerged in evolution over 400 million years ago in bony fish and throughout phylogeny they have evolved to ensure our survival. They control the volume and composition of our extracellular fluid, maintain normal arterial pressure, tissue perfusion, and oxygen delivery to vital tissues. During early mammalian development, the cells are widely distributed throughout the kidney vasculature where they are crucially involved in the assembly and morphogenesis of the kidney arteries and arterioles^1^. In the adult animal, they retain some progenitor properties as they can regenerate damaged glomeruli, the filtering units of the kidneys^2,3^. As a result of the aforementioned morphogenic functions, in early embryonic life renin cells are broadly distributed throughout the renal vasculature, and as the cells differentiate, adult renin cells are confined to the tip of the renal arterioles near the glomerulus, thus their name juxtaglomerular (JG) cells^4–7^.

Whereas significant work has uncovered several pathways and genomic regions integral to the operation of renin expressing cells, understanding the epigenetic changes that occur to regulate the renin phenotype is a major goal of several laboratories around the world. Using lineage tracing techniques, we and others have shown that all the mural cells of the kidney arteries and arterioles, including renin expressing cells, descend from *FoxD1*+ stromal progenitors. Yet, the factors and mechanisms that control the differentiation of these early progenitors into intermediate cells and ultimately renin cells, vascular smooth muscle cells (VSMCs), and mesangial cells (MCs) remains poorly understood. We know that the renin locus contains a cAMP responsive element (CRE) where the histone acetyl transferases CBP/p300 can bind to regulate renin expression^8–13^. Phosphorylation of the transcription factor (TF) Creb and binding to the CRE located in the 5’ regulatory region of the renin gene activates renin gene transcription. In addition, the final common effector of the Notch signaling pathway, RBP-J (Recombination signal binding protein for immunoglobulin kappa J region), is necessary to maintain renin expression and modulates the plasticity of VSMCs and MCs to re-acquire the renin phenotype when homeostasis is threatened^6,14–17^. RBP-J also regulates *Akr1b7* (Aldo-keto reductase family 1, member 7) which is co-expressed with renin and serves as an additional marker of mature renin cells^6,18^. Past work in our group identified a set of super-enhancers unique to renin cells^7^. The primary super-enhancer was found just upstream of the renin gene (*Ren1*) and is thought to harbor the memory of the renin phenotype in renin cell descendants^7^.

While these efforts have greatly contributed to our knowledge about promoter and enhancer elements affecting renin expression^10,13,15,19–23^, no comprehensive study has been performed to identify the chromatin states, genomic loci, and specific transcription factors (TFs) that determine the differentiation trajectory of the renin cells from their early, stromal *FoxD1* progenitors to the mature JG cell. To identify those landmark events, we constructed a single-cell atlas of chromatin accessibility and gene expression profiles along a comprehensive developmental time course of renin cell development and other *FoxD1* descendants and identified the critical regulators that determine their identity and fate.

## Methods

Comprehensive materials and methods information is available in the Supplemental Material.

### Data and Code Availability

- The single cell omics data have been uploaded to the Gene Expression Omnibus (GEO) with the accession IDs GEO: GSE218570.
- Any additional information is available from the Lead Contact(s), R. Ariel Gomez, upon request.

## Results

To investigate the epigenomic and transcriptomic changes that occur during kidney vascular development, we isolated single cells from *FoxD1cre;mTmG* mice across four developmental time points at embryonic day (E) 12, E18, post-natal day (P) 5, and P30. By P30, renin expressing cells are confined to the JG region, MCs are localized within the glomeruli, VSMCs constitute the walls of the renal arterioles, and pericytes (PCs) surround them. All of the *FoxD1* lineage cells retain the memory to revert to renin expressing phenotypes when homeostasis is threatened. In our lineage-tracing model, cells that express *FoxD1* at any point during development were enduringly labeled by the expression of GFP. Isolated cells were subjected to scATAC-seq and scRNA-seq experiments. We identified the developmental trajectory that gives rise to the rare endocrine JG cell population using *FoxD1* as a marker of cells that initiate the trajectory. In turn, JG cells were identified by the concomitant expression and accessibility at *Ren1* and *Akr1b7*. We investigated epigenetic and transcriptomic changes independently at each time point, and identified the rare cell population of renin expressing cells.

### Time-point analysis unravels cell fate decisions in the assembly of the kidney vasculature

To investigate the cells responsible for forming the kidney vasculature, we analyzed *FoxD1*+ cells at E12. Based on the top gene markers derived from the scRNA-seq data, we annotated E12 cells as: “Nephron Lineage”, “Proliferating Cells”, “Nephrogenic Zone Stromal Cells”, “Stromal Cells”, “Mesenchymal/Stromal Cells”, “Committing Nephron Progenitor”, and “Ureteric Epithelium” (Fig. 1A). At E12, differentiation is not uniform with few differentiated cell populations present and labeled as “Committing Nephron Progenitors” and “Nephron Lineage” cells. These two clusters express the classical marker for the posterior intermediate mesoderm, the Odd Skipped Related Transcription Factor 1 (*Osr1*). As the differentiation pathway proceeds from the intermediate mesoderm to the loose metanephric mesenchyme, cells begin expressing *FoxD1* as the *Osr1* expression decreases (Fig. 1B). Indeed, the variance of the *FoxD1* gene expression level is high (Fig. 1C, Fig. S1A), confirming this process is not uniform. The “Proliferating Cells” cluster had high *FoxD1* expression, but insufficient additional marker expression to be annotated further. Additionally, some *FoxD1*+ cells expressed markers from other embryonic kidney structures, such as ureteric bud (marked by *Hoxb7*) (Fig. S1B) or the cap mesenchyme (marked by *Six2* (Fig. S1C) and *Cited1* (Fig. S1D)). This overlap is in accordance with our previous histological findings^24^. Renin starts to be expressed in the kidney around E12^23^, yet at this stage we found few renin expressing cells (Fig. S1E, F) or VSMCs (Fig. S1G, H). Nevertheless, two clusters showed the highest *FoxD1* gene expression levels, the “Nephrogenic Zone Stromal Cells” and “Stromal Cells” (Fig. 1C). Evaluating clusters with high *FoxD1* expression revealed highly enriched motifs for the TFs *Hoxb9, Hoxd13, Hoxd10*, and *Hoxa10*, which may drive Sonic hedgehog (*Shh*) expression to orient cells along the anterior to posterior axis^25^. We next integrated the scRNA-seq data with the scATAC-seq data and performed motif enrichment analysis (Fig. 1D). Within the “Stromal Cells” cluster, in which *FoxD1* gene expression levels are the highest, we also identified enriched motifs for *Tcf21* and more (Fig. 1D). *Tcf21* is enriched during early nephrogenesis and is essential for kidney development. *Tcf21* has pleiotropic functions during nephrogenesis and is required for the normal crosstalk among the loose mesenchyme, cap mesenchyme and ureteric bud which ultimately leads to the branching morphogenesis^26^. Using RNAscope^27^, we confirmed *Tcf21* expression early throughout the developing kidney, suggesting it is an important factor for *FoxD1* progenitor differentiation at E12, although *Tcf21* does not co-localize specifically with renin-expressing cells and decreases in expression as development progresses (Fig. S2).

**Fig. 1:**
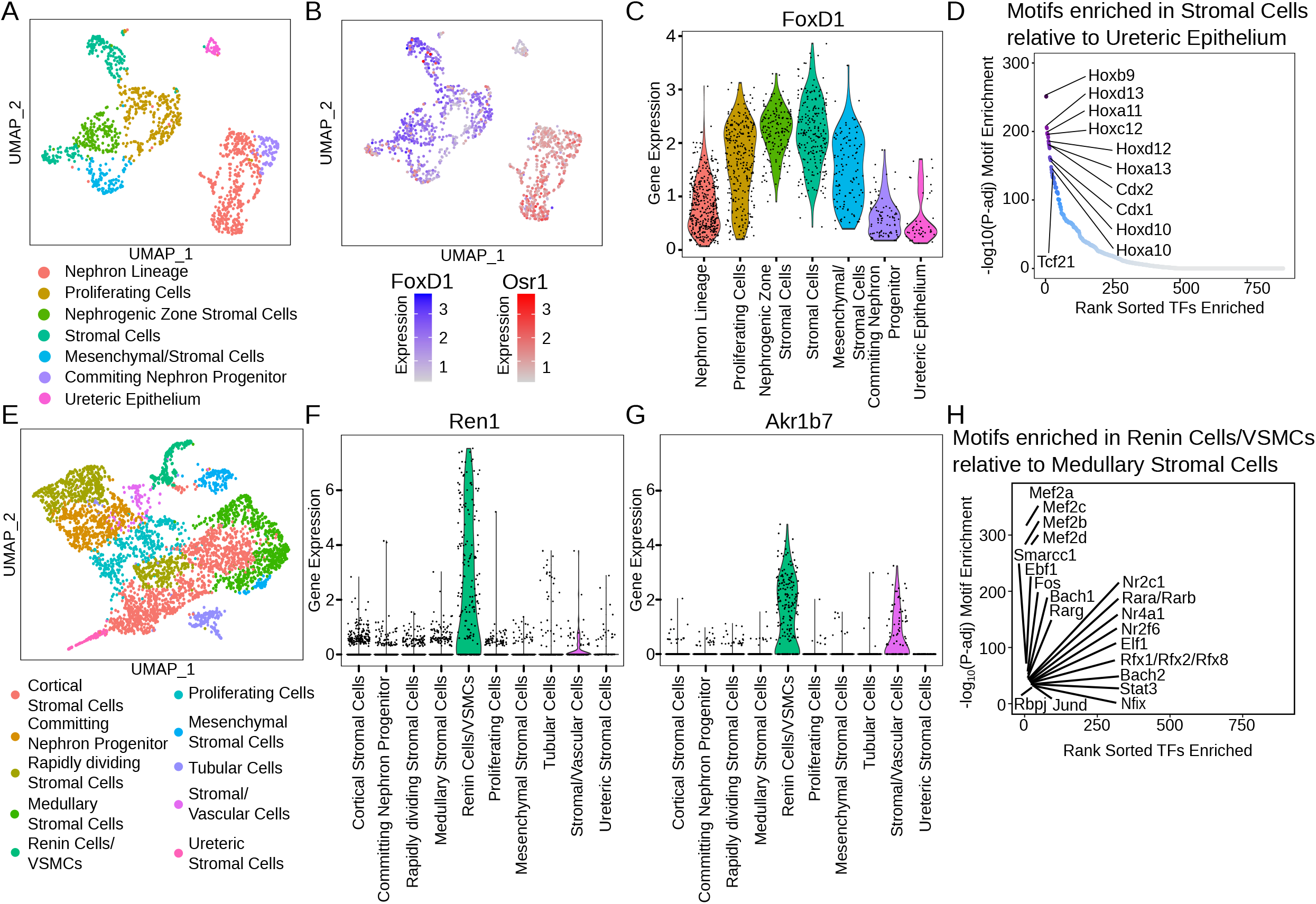
FoxD1+ cell differentiation reveals cells committed to the nephron lineage. (A-D) E12. (A) UMAP visualization of annotated scRNA-seq data. There are clusters committed to nephron progenitors and lineage members with a subset of cells already differentiated into stromal cells. (B) UMAP visualization of the gene expression distribution for Osr1 and FoxD1. The differentiation pathway from Osr1 towards the FoxD1 clusters is easily detectable from right to left. (C) FoxD1 gene expression among the identified cell populations. The “Stromal Cells” and “Nephrogenic Zone Stromal Cells” clusters have the highest FoxD1 expression. These cells are poised to differentiate into kidney arteries and arterioles cell populations.Top enriched TF motifs in “Stromal Cells” relative to the “Ureteric Epithelium”. (E-H) E18. (E) UMAP of FoxD1 derivative cells with their respective annotations. (F) Renin gene expression in all clusters. Although there is high variability, the “Renin Cells/VSMCs” cluster has the highest expression. (G) Violin plot showing Akr1b7 gene expression in all clusters. The “Renin Cells/VSMCs” cluster has the highest overall expression of Akr1b7. (H) Top enriched TFs in the “Renin Cells/VSMCs” cluster detected within the differential accessible peaks compared against “Medullary Stromal Cells”.

At E18, mice gestation is nearing completion (E19-E20), but nephrogenesis continues 5-7 days after birth. *FoxD1* derivatives are still actively differentiating and proliferating, but we identified cell populations characteristic of later developmental stages (Fig. 1E). The “Stromal Cells” can be distinguished by kidney spatial region such as cortical and medullary stromal cells. Here, we observe two cell clusters directly related to the vasculature: “Stromal/Vascular Cells” and “Renin Cells/VSMCs”. Although no single cluster was assigned as pericytes, both of the aforementioned clusters have markers related to them, such as Melanoma Cell Adhesion Molecule (*Mcam*) (Fig. S1I), Chondroitin Sulfate Proteoglycan 4 (*Cspg4*) or Neuron-glial antigen 2 (*Ng2*) (Fig. S1J), and Nestin (*Nes*) [30] (Fig. S1K). Pericytes contained within those clusters explain the observation that in the fetus or adult animal subjected to a homeostatic threat, pericytes synthesize renin^28^.

As expected, the “Renin Cells/VSMCs” cluster displays the highest *Ren1* and *Akr1b7* expression (Fig. 1F, G). The renin gene is a *sine qua non* marker for the renin progenitor cell (RPC), while *Akr1b7* was identified more recently in our lab as an independent marker of renin cells^14,18^. The presence of cells with high levels of both *Ren1* and *Akr1b7* suggests some cells have already acquired their definitive fate, becoming JG cells. The “Renin Cells/VSMCs” cluster also expressed high levels of smooth muscle genes such as Smooth Muscle Alpha Actin (*Acta2*) (Fig. S1L), Smooth Muscle Myosin Heavy Chain 11 (*Myh11*) (Fig. S1M), Transgelin (*Tagln*) (Fig. S1N), and Calponin 1 (Cnn1) (Fig. S1O). A large body of evidence support the RPCs as the main precursors of the majority of mural cells of the kidney arteries and arterioles, including the VSMCs^15,29,30^. Therefore, the VSMCs gradually acquire the smooth muscle transcriptome while renin and *Akr1b7* gene expression remains highest in what ultimately form the JG cell population.

*Rbpj*, which is enriched in renin expressing cells present at E18 (Fig. <u>1</u>H), is an important regulator of the renin cell phenotype with *Rbpj* deletion causing renin cells to stop making renin and/or smooth muscle proteins, instead adopting an abnormal “hematological” phenotype^17^. It seems that *Rbpj* plays a dual role in renin cells: 1) it orchestrates a group of genes that determine the myo-endocrine renin cell phenotype, and 2) prevents cells from the undesirable ectopic expression of genes from non-renin cell lineages^17^. The Fos family heterodimerizes with the Jun family to form the Activator protein 1 (*AP-1*) TF, which regulates differentiation, proliferation, apoptosis, and senescence^31,32^ and members of both families are enriched among renin expressing cells and VSMCs at E18 (Fig. 1H). Our lab identified a set of super-enhancers that regulates *Ren1*^7^. Among them, *Junb* is the third ranked enhancer, suggesting its strong participation in the renin cells biology. We are currently deleting *Junb* in the *FoxD1* lineage to determine whether it affects renin cell identity.

Within the “Renin Cells/VSMCs” population at E18, the myocyte enhancer factor 2 (MEF2) family (*Mef2a, Mef2c, Mef2b, Mef2d*) of TFs is enriched, highlighting the significance of this TF family to the “Renin Cells/VSMCs” differentiation (Fig. 1H). We confirmed the presence of MEF2 members in the kidney and identified *Mef2b* in particular as co-localizing with renin expression as early as E18 (Fig. S2). Furthermore, since this cluster may include JG cells, RPCs, VSMCs, and PCs the enrichment of MEF2 members strongly suggest a pivotal role for this TF family within *FoxD1* derivative cells.

At P5, nephrogenesis is nearly complete. The majority of the *FoxD1* derivatives cells are terminally differentiated (Fig. S3A). We annotated clusters of “Mesangial Cells”, “Fibroblasts”, “Podocytes”, “Renin Cells/VSMCs”, and “Immune Cells”. Although podocytes are not derived from *FoxD1* stromal cells, it is known these cells express *FoxD1* after birth^33^. Unfortunately, we were not able to identify a cluster specific for JG cells within the independent timepoint analysis, but one cluster was assigned as “Renin Cells/VSMCs”. Interestingly, *Ren1* and *Akr1b7* both display high variance between clusters (Fig. S3B, C). The data suggest that low renin expressing cells are terminating their differentiation, while cells with concomitant high expression levels of renin and *Akr1b7* fully differentiate into JG cells. Motif enrichment analysis identified the MEF2 family as highly enriched in the “Mesangial Cells” cluster (Fig. S3D), as were the TFs *Elf1, Ebf1, Elf3*, and *Elf5*. The “Renin Cells/VSMCs” cluster again exhibited the MEF2 family as the highest enriched, although other TFs such as *Nfix, Nfic, Smarcc1, Bach1, Hic1*, and *Hic2* were also identified.

At P30, all the vasculature has been assembled and under basal conditions without homeostatic stressors, all the renin expressing cells are now JG cells^4^. Clustering assignments were further refined at this stage (Fig. S3F), although specific assignment of a single cluster to exclusively JG cells remained challenging. Nevertheless, *Ren1* gene expression variance in the renin expressing cell clusters remained high (Fig. S3G, H). This suggests RPCs have fully completed their differentiation pathway and only the JG cells endure with high renin and *Akr1b7* expression levels. Pairwise motif enrichment analysis revealed, once again, the MEF2 family as important drivers of the joint “Renin Cells/VSMCs” cluster along with *Nfix, Nfic, Stat3*, and *Stat4* (Fig. S3I). Taken together, the data suggests that MEF2 family members are major drivers of the *FoxD1* precursor’s differentiation trajectory.

### Epigenetic landscape of renin cell development

A major motivation for this study was the identification of the mature renin expressing JG cells in the mouse kidney, and the discovery of the epigenomic and transcriptomic changes and features unique to this cell population. Whereas independent investigation of each timepoint uncovered features unique to broadly defined renin expressing cell populations, we next sought to integrate data across development to construct a developmental trajectory that pinpoints the JG cell populations.

The scATAC-seq samples formed clusters predominantly separated by developmental timepoint with P30 cells showing the most spatially removed clustering profile (Fig. 2A). Independent scRNA-seq cells were clustered and revealed 22 distinct annotated clusters (Fig. 2B). We integrated the scRNA-seq data with the scATAC-seq and performed label-transfer to ultimately annotate 20 open chromatin derived clusters (Fig. 2C).

**Fig. 2:**
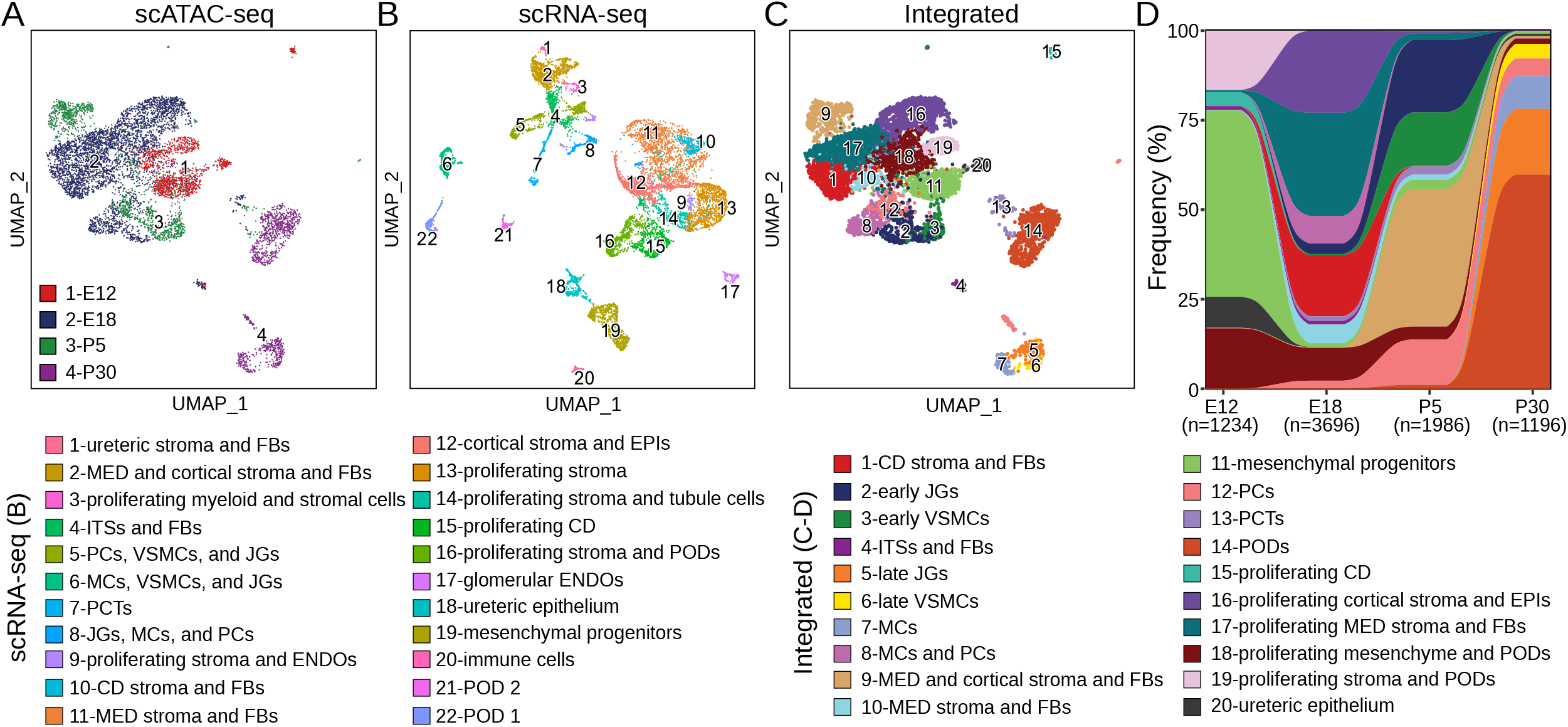
Overview of scATAC-seq, scRNA-seq, and integrated data visualizations identifying the rare renin cell (JG) populations. (A) UMAP visualization of scATAC-seq data colored by developmental timepoint. (B) UMAP visualization of scRNA-seq data with cell annotations across development. (C) UMAP visualization of integrated scATAC-seq and scRNA-seq data with cell annotations following identification of the JG population. (D) Alluvial plot of cell frequency distribution across developmental time point. Numbers below time points represent total number of single cells at each time point. (CD: collecting duct; ENDOs: endothelial cells; EPIs: epithelial cells; ITSs: interstitial cells; JGs: juxtaglomerular cells; MCs: mesangial cells; MED: medullary; FBs: fibroblasts; PCs: pericytes; PCTs: proximal convoluted tubules; PODs: podocytes; VSMCs: vascular smooth muscle cells)

We also explored the distribution of annotated cells in the open chromatin data across developmental time points to evaluate lineage contributions to the JG cells (Fig. 2D). To identify the subset of cells representing the JG population, we looked for canonical markers of JG cell identity, the genes *Ren1* and *Akr1b7*^23^. By evaluating these marker genes from both gene activity scores and integrated gene expression (Fig. S4B, C), we identified the subpopulation of JG cells.

### Differentiation trajectory of JG cells

Much remains unknown about the epigenetic changes that lead to JG cell identity. To resolve this, we sought to identify the regions of open chromatin, corresponding TFs, and their linked gene score and expression to investigate what genetic and epigenetic features define JG cells. Because mature renin cells are so rare, only a handful of studies have successfully identified markers for these cells using cell labeling followed by sorting and microarray analysis^14^, or by isolating a large number of unlabeled cells^34^. Our study design specifically enriches for cells in this lineage so that even with a limited number of total cells we identified this rare population. We defined a pseudo-time trajectory of cells with high *FoxD1* expression and accessibility beginning in E12 through E18 to cells with high expression and accessibility of *Ren1* and *Akr1b7* to identify the JG cell population, which was apparent as early as E18 and present at P5 and P30 (Fig. 3A, Fig. S4A-C, G). We then evaluated the expression and accessibility of VSMC markers, *Cnn1, Acta2*, and *Myh11*, to determine where along the trajectory VSMCs appear and differentiate from JG cells (Fig. S4D-G).

By looking within peaks that mark individual cell clusters along the developmental trajectory, we identified differentially enriched motifs in individual cell populations (Fig. 3B, C). We identified significant enrichment of the MEF2 family of TFs and *Nfix, Ebf1, Zfa*, and *Sp2* in JG cells and to a lesser extent in VSMCs (Fig. 3B, C; Fig. S5A, B, J). MEF2 family enrichment occurs during the formation of the early JG cell population and remains enriched throughout JG cell differentiation. The nuclear factor I (NFI) family of TFs (including *Nfix* and *Nfic*) have been previously reported to bind to promoter and enhancer elements at the *Ren1* locus^20,21^, and we confirm motif occurrences of these and other factors at the *Ren1* locus (Fig. S4I-J). In the adult kidney, *Nfix* is found in the proximal tubules^35^ and plays a pivotal role in extracellular matrix (ECM) generation. Some studies have suggested that *Nfix* would be a target to ameliorate muscular dystrophies^36^, and it has an epigenetic participation in neural stem cell quiescence^37^. Whether *Nfix* affects renin cells, VSMCs and/or pericytes during chronic stimulation of the RAS and contributes to concentric vascular hypertrophy requires further investigation. To confirm the role of *Nfix* in JG cell development, we first evaluated the presence of *Nfix* by RNAscope (Fig. S2) and found that *Nfix* exhibited some co-localization with renin by E18 through P30. At higher magnification we see *Nfix* wrapping renin-expressing cells, suggesting its expression at PCs (Fig. S2). Our past bulk RNA-seq data showed that *Nfix* is the highest expressed TF in renin producing As4.1 cells with a TPM value of 277. Therefore, we sought to determine whether *Nfix* plays a role in the regulation of *Ren1* expression. We knocked out *Nfix* in As4.1 cells using CRISPR-Cas9 (Fig. 4). We targeted three sgRNAs to exon 2 of *Nfix*, an exon present in all protein encoding variants of the gene (Fig. 4A). Using a GFP reporter plasmid, the efficiency of As4.1 cell transfection after nucleofection was 92% (Fig. 4B) and, we obtained a highly efficient knockout (99%) of *Nfix* as determined by Sanger sequencing and ICE analysis (Fig. 4C, D). qRT-PCR analysis showed a significant decrease of *Ren1* mRNA levels in *Nfix* KO cells compared to controls (Fig. 4E. Control: 1.014 ± 0.197; *Nfix* KO: 0.004 ± 0.0003, n=3, p=0.009, two-tailed t test). This demonstrates that *Nfix* plays an essential role in the control of *Ren1* expression in cultured renin cells and agrees with a previous study by Pan et al. 2003^38^.

**Fig. 3:**
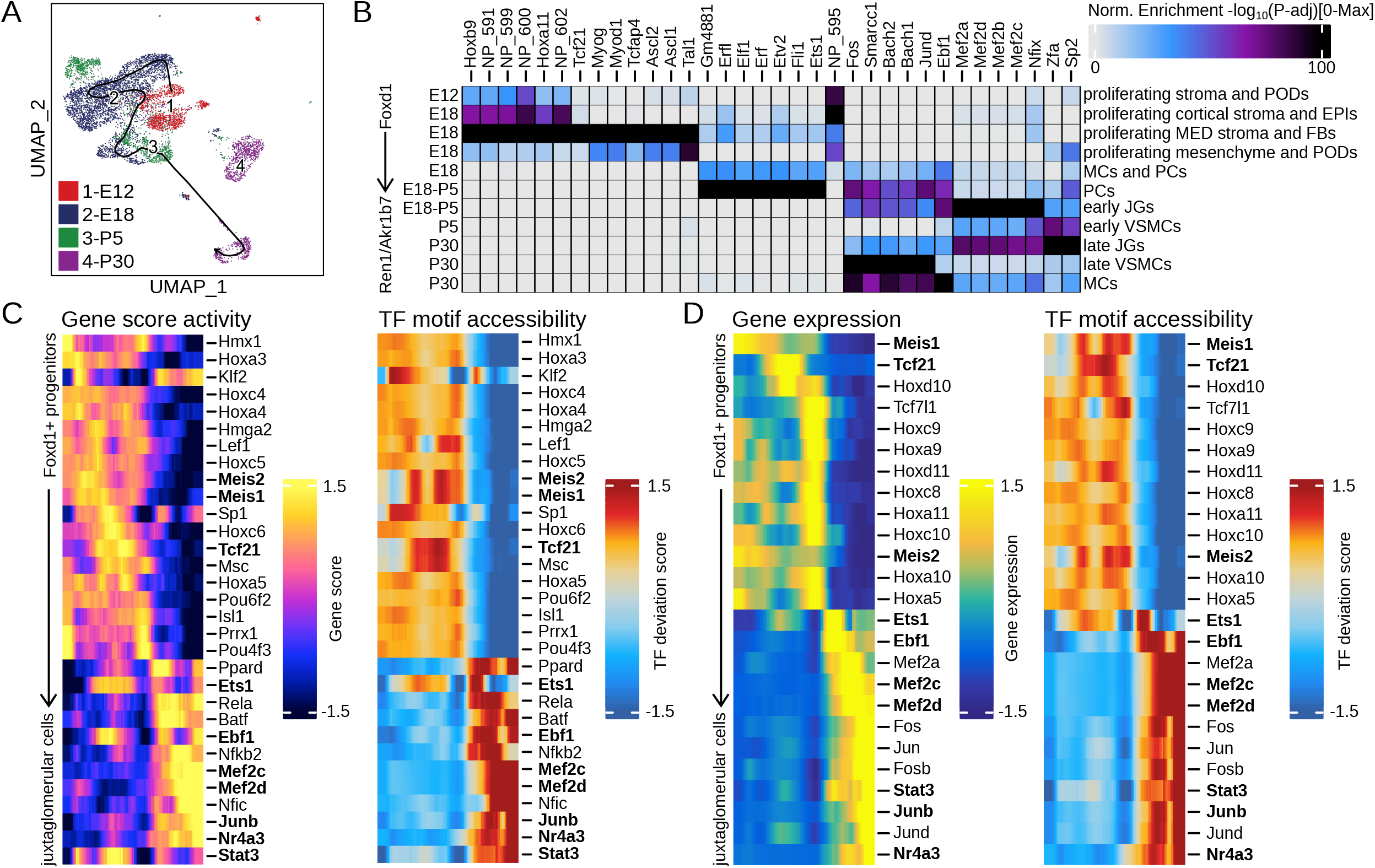
Epigenomic differentiation trajectory identifies renin cells by embryonic day 18 in mouse kidney development. (A) UMAP visualization showing the pseudo-time trajectory across developmental time points. Arrow head represents the end point of the trajectory. (B) Heatmap of motif hypergeometric enrichment-adjusted P values within the marker peaks of each JG trajectory cluster. Color indicates the motif enrichment (-log(sub)10(/sub)(P value)) based on the hypergeometric test. Left y-axis labels timepoint of clusters identified along the right y-axis. (C) Integrated pseudo-time analysis of positively correlated gene scores and motif z-scores. (D) Integrated pseudo-time analysis of positively correlated gene expression and motif z-scores. **Bold** text indicates genes/motifs identified across both approaches. (AA: afferent arteriole; EA: efferent arteriole; EPIs: epithelial cells; JGs: juxtaglomerular cells; MCs: mesangial cells; MED: medullary; FBs: fibroblasts; PCs: pericytes; PODs: podocytes; VSMCs: vascular smooth muscle cells)

**Fig. 4:**
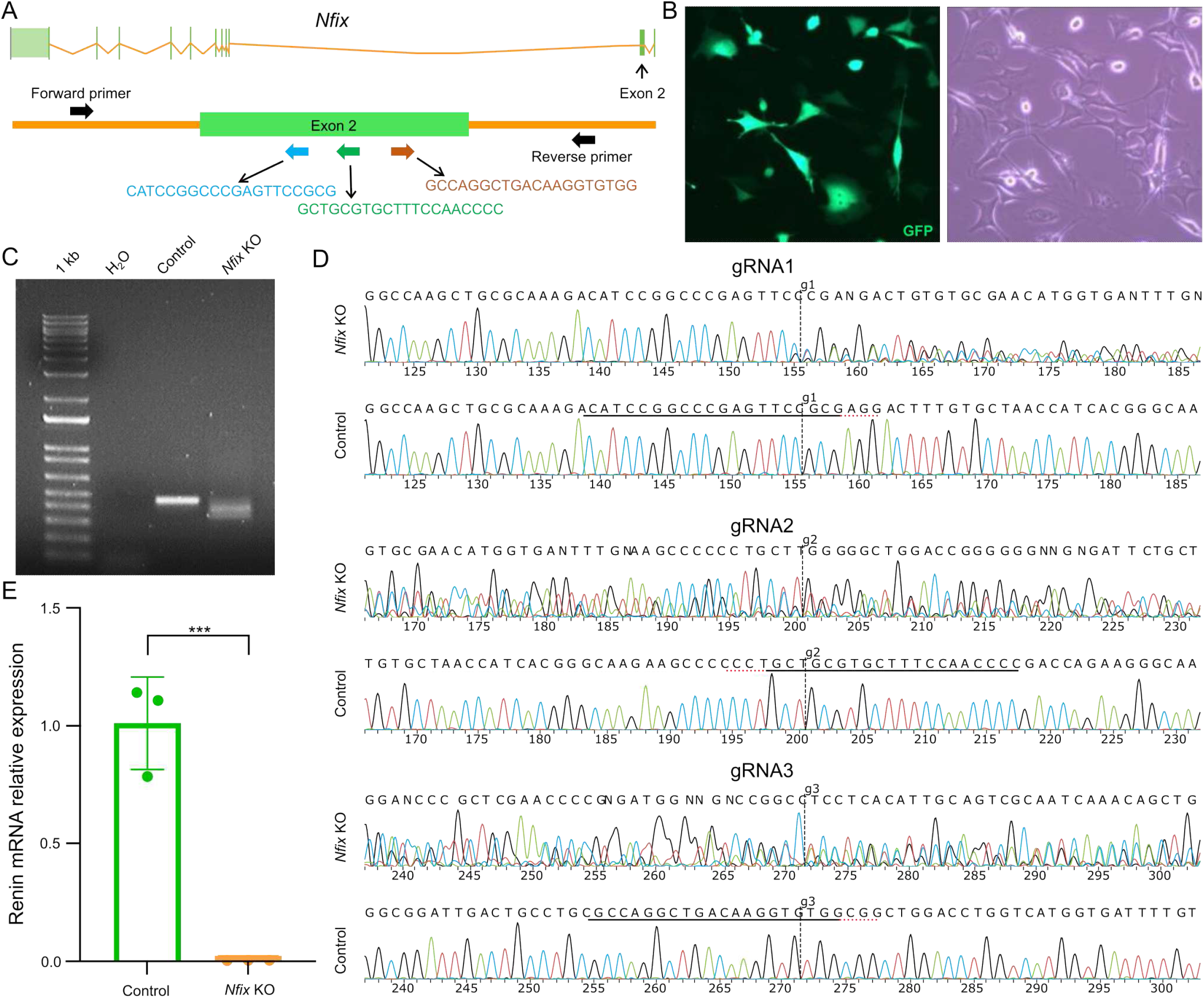
Deletion of Nfix in cultured renin cells decreases Ren1 mRNA expression. (A) Schematic showing the guide RNA (gRNA) targeting sites on exon 2 of the Nfix gene. Depicted is transcript Nfix-204 (ENSMUST00000109764.8) which has 11 exons. Three gRNAs were designed for Nfix knockout (KO). Primers designed to confirm Nfix KO are indicated by black arrows. (B) Images of AS4.1 cells 48h after nucleofection with pmaxGFP plasmid. (Left panel: GFP image; right panel: phase image.) The efficiency of transfection was 92%. (C) Agarose gel electrophoresis of the PCR product using genomic DNA isolated from Nfix KO renin cells showed three DNA fragments compared to one 448bp product in control cells. (D) Analysis of CRISPR-mediated mutations by Sanger sequencing. Shown are the edited and wild-type (control) sequences in the region around the three guide sequences. The traces show sequence base calls from both the control and the experimental sample .ab1 files, which contain mixed base calls. The guide sequences are underlined in black and the PAM sequences in red dashed lines. The vertical black dotted lines represent the cut sites. Mixed sequencing bases after the cut indicate that cutting and error-prone repair occurred. (E) Quantitative reverse transcription PCR showed a significant decrease in Ren1 mRNA in Nfix KO cells (n=3, p=0.009, two-tailed Student’s t-test). Data are means ± standard deviation.

Renin progenitors have been identified in mouse bone marrow that express renin and *Ebf1*, a key pioneer TF of B-cell specification and commitment^39^, and they are the cell of origin of a highly penetrant B-cell leukaemia^40^. Thus, hematopoietic and endocrine cells that express renin have in common the utilization of *Ebf1* in a manner that is intriguing and remains to be explored. Whether *Ebf1* plays a role in the identity of the renin cell as it does in B lymphocytes remains to be determined.

We also utilized our integrated pseudo-time analysis to identify positive drivers of differentiation along the trajectory (Fig. S4H). We correlated gene scores or gene expression to their corresponding motifs and uncovered numerous linked factors including the MEF2 family, *Ebf1, Junb, Nr4a3*, and *Stat3* as enriched across both integrative approaches in JG cells (Fig. 3D, E). We also identified gene scores, gene expression, regulatory regions, and TF motifs across pseudo-time that are enriched but not positively correlated.

By the emergence of early (E18-P5) and late JG cells (P30), there are enriched gene scores for *Lama5, Rrad, Hspb1, Arid5a, Gja4*, and *Pdgf-a* (Fig. S5K). *Lama5*, which produces laminin-*α*5, is an essential component of the glomerular basement membrane and mutations in this gene are associated with chronic kidney disease^41,42^. Past work has stressed the importance of lamin A/C for sensing and responding to extracellular physical forces to regulate renin expression^43^, and *Lama5*, although notably different from lamins, offers an intriguing avenue for future study as a potential coplayer in the extracellular matrix to transduce pressure or mechanical signals to renin expressing cells. *Rrad* is a calcium channel regulator previously reported as associated with prorenin^44^, and it is known that renin secretion from JG cells is regulated via calcium signaling pathways^45,46^. *Rrad* may represent an understudied factor regulating JG cell development. *Hspb1* is an anti-apoptotic protein that is itself increased by the presence of angiotensin II^47^. It is known to be upregulated in diabetic nephropathy, typically in podocytes^47^, and as an apoptotic inhibitor following acute kidney injury in renal tubular cells^48^. Here, the data is suggestive of a possible role in the initial diferentiation of JG cells. *Arid5a* as an RNA-binding protein which has generally been understood to respond to inflammation. It is known to translocate the cytoplasm and stabilize *Stat3*, which we identify as also enriched in JG cell populations^49^. *Arid5a* is activated by NF-*κ*B^50^, which itself can induce upregulation of the prorenin receptor (PRR)^51^, although the affinity of renin for this receptor is understood to be quite low^52^. We are currently deleting this receptor in a renin cell model (PRR^*fl/fl Ren1dCre*; data not shown), which interrupts the ability of VSMCs to be transformed into the renin phenotype, suggesting that the PRR is essential for renin bioavailability. *Gja4*, also known as connexin 37, has been previously reported as expressed in VSMCs of hypertensive animals and in renin-secreting cells^53^. Previous studies of renin related genes demonstrated the importance of the connexin family, particularly connexin 40^14^ in renin expression. Connexin 37 is reported to play a role in angiotensin II signaling through the modulation of the angiotensin II type 2 receptor^53,54^. Here, we indicate a possible role for *Gja4* as a positive driver of differentiation of mature JG cells. The PDGF family is well studied in renal fibrosis, and *Pdgf-a*, which is significantly enriched in late JG populations, has previously been found upregulated in mesangial, endothelial, and smooth muscle cells of the fibrotic kidney^55,56^.

The JG marker genes *Ren1* and *Akr1b7* and VSMC marker genes *Acta2, Tagln, Crip1*, and *Myh11* are enriched as expected (Fig. S5L). We further visualized the enrichment of JG defining lineage markers by showing that *FoxD1* is expressed and accessible early along the trajectory with *Ren1, Akr1b7*, and VSMC marker genes increasing in expression and accessibility in their expected cell populations (Fig. S4G).

When JG cells have emerged, there are enriched regulatory regions in chromosomes 1, 2, 4, 6, 7, and 8 (Fig. S5M) indicating possible trajectory defining regulatory regions. Overall, we again identified enrichmed motifs for the MEF2 family of TFs, as well as motifs for *Smarcc1, Bach1* and *Bach2, Nfix, Fos, Fosb, Hic1* and *Hic2, Snai2*, and *Jund* (Fig. S5I, N).

*Smarcc1* is a chromatin remodeling enzyme, whose enzymatic activity changes the chromatin structure by altering DNA-histone contacts within a nucleosome in an ATP-dependent manner^35^. We classified *Smarcc1* as an early TF that comes in to play early during renin cell differentiation, potentially by eliciting a favorable chromatin landscape for renin gene expression.

*Bach1* and its paralog *Bach2* are involved in transcriptional activation or repression via *Mafk* (bZip Maf transcription factor protein) and form heterodimers with small Maf proteins which bind at Maf-recognition elements (MARE) of target genes^57,58^. MAREs share strong sequence conservation with CRE elements^59^ with a cAMP response element present at the renin locus and essential for renin expression^6,7,60^. The Maf family of TFs are important regulators of kidney development and differentiation and are implicated in both normal development and pathophysiological processes responsible for a variety of congenital kidney diseases. For instance, *Mafb* homozygous mutation displayed renal dysgenesis with abnormal podocyte differentiation, fusion and effacement of pedicels, and tubular apoptosis leading to congenital nephrotic syndrome^61^. Moreover, *Mafb* mutations in humans result in focal segmental glomerulosclerosis with Duane Retraction syndrome^62^. Whether these TFs can interfere directly or indirectly with the renin cells’ physiology requires further investigation.

### Transcription factors contributing to JG cell development

We observed a pattern of expression and accessibility changes that categorized specific TFs as important during different stages of JG cell development. In addition to *Smarcc1* as mentioned above, we find significant enrichment of the NFI TF family member, *Nfib* (Fig. 5A), in the late differentiation clusters that include the JG cell populations. *Nfib* gene expression (Fig. 5E) and activity score (Fig. 5I) peaks early along the JG differentiation trajectory before progressively lessening over time, although both expression and accessibility rise during the formation of late forming JG cells. Enriched chromVAR deviation scores across the trajectory confirm the significance of *Nfib* in the differentiation of these cells. (Fig. 5M). *Nfib* is related to the progression of renal clear cell carcinomas^63^, and the related *Nfia* is associated with abnormalities of the ureteropelvic and ureterovesical junctions^64^. Further studies will be required to define the role of *Nfib* in renin cell identity.

**Fig. 5:**
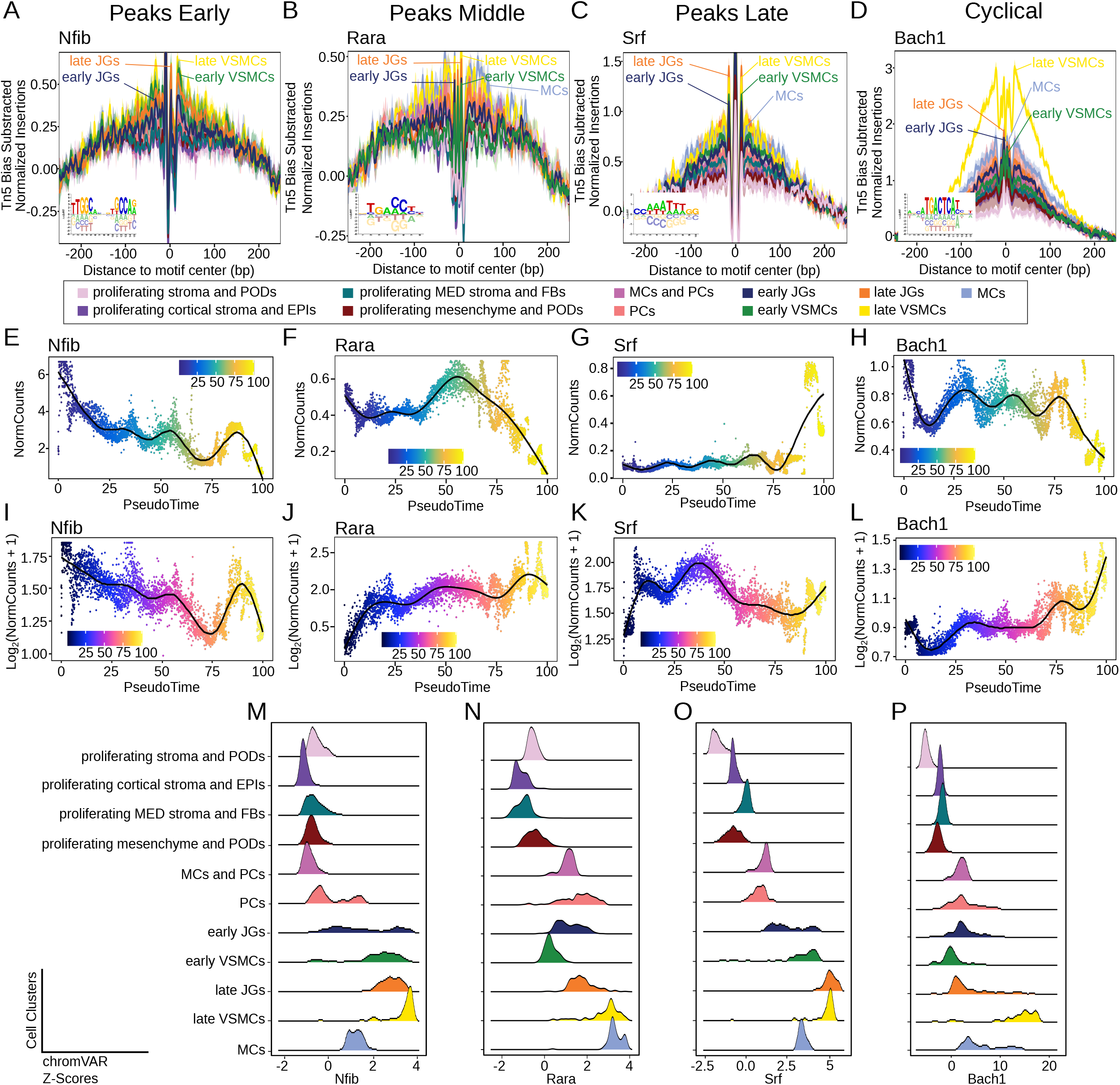
TF expression, accessibility, and enrichment uncover patterns of differentiation. Tn5 insertion bias corrected footprints for Nfib (A), Rara (B), Srf (C), and Bach1 (D) are enriched in JG and VSMC clusters (Inset represents the corresponding motif logo). The expression pattern of enriched TFs in the JG cluster illustrates early (E), middle (F), late (G), and cyclical (H) patterns of transcript abundance. Gene score activity generally recapitulates expression patterns of early (E), middle (F), late (G), and cyclical (H) activity. TF chromVAR deviation z-scores highlight enrichment of Nfib (m), Rara (N), Srf (o), and Bach1 (P) in middle to late differentiation along the JG trajectory. (EPIs: epithelial cells; JGs: juxtaglomerular cells; MCs: mesangial cells; MED: medullary; FBs: fibroblasts; PCs: pericytes; PODs: podocytes; VSMCs: vascular smooth muscle cells)

We identified enriched motifs for members of the retinoic acid receptor (RAR) family of TFs with an enriched footprint for *Rara* specifically in JG and late forming cell populations (Fig. 5B). *Rara* displays a pattern of increased transcript abundance along the middle of the trajectory with an enrichment in accessibility late among clusters forming late JG and VSMCs (Fig. 5F, J, N). A second RAR family member, *Rarg*, is also enriched during the initial differentiation of JG cells with an overall enrichment in late differentiation clusters (Fig. S5C).

*Srf* also displayed an enriched footprint in both JG and VSMC populations with comparatively high transcript abundance and an enriched chromVAR deviation score in these late cell populations (Fig. 5C, G, K, O). Interestingly, *Bach1* undergoes a cyclical pattern of gene expression (Fig. 5H) and an enrichment in accessibility late (Fig. 5L) emphasizing a possible role in the cell cycle of differentiating JG cells^65,66^. Each of the aforementioned TF motifs are preferentially enriched in late forming cell clusters, including the early and late JG populations (Fig. 5M-P).

In finding that a number of TFs were important at different stages of JG cell differentiation, we categorized additional TFs as enriched early, middle, or late during JG cell differentiation (Fig. S6). TFs important early in differentiation included *Arid3a, Tcf21, Cdx1, Hoxa9*, and *Setbp1* (Fig. S6A). As discussed above in relation to *Arid5a*, Arid family members are involved in cell differentiation and proliferation and are emerging targets in numerous cancers^67–69^. *Arid3a*, in particular, has been associated with nephric tubule regeneration^70^ and its relevance to JG cell differentiation and the ability of JG cells to regenerate damaged glomeruli^22,23,71^ supports a signficant role for this protein. TFs enriched along the middle of the JG cell trajectory included *E2f1, Egr1, Foxn3, Sp1*, and *Rbpj* (Fig. S6B). E2F TFs also play important roles in the cell cycle, and *E2f1* is particularly important in MC proliferation^72^. Previous studies^20,21^ have shown that Sp TFs, including *Sp1*, can bind at renin regulatory regions, which we confirmed bioinformatically (Fig. S4I). In later differentiation, we identified *Ahctf1, Creb1, Lcor, Pgr*, and *Zfa* as enriched in JG cell and VSMC populations (Fig. S6C).

### MEF2 family of TFs are uniquely enriched in JG cells

As *Ren1* activity is paramount to JG cell function and identity, we predicted regulators of cells with or without renin expression to identify major drivers of this differential activity. Using an adaptive elastic net implemented in `RENIN`^73^, we predicted which regulatory elements and TFs are likely to regulate *Ren1*. MEF2 family members are predicted as top regulators of renin activity (Fig. 6A). For greater nuance of JG cell identity, we investigated which factors specifically differentiated early JGs from the nearest trajectory population of PCs. We corroborated the enrichment of MEF2 members in JG cells (Fig. 6B). Futher supporting the role of *Nfix* and other NFI members, both NFI and regulatory factor X (RFX) TF families motifs are enriched in JG cells with corresponding binding sites at both *Ren1* and *Akr1b7* within JG marker regions (Fig. 6B, I-J).

**Fig. 6:**
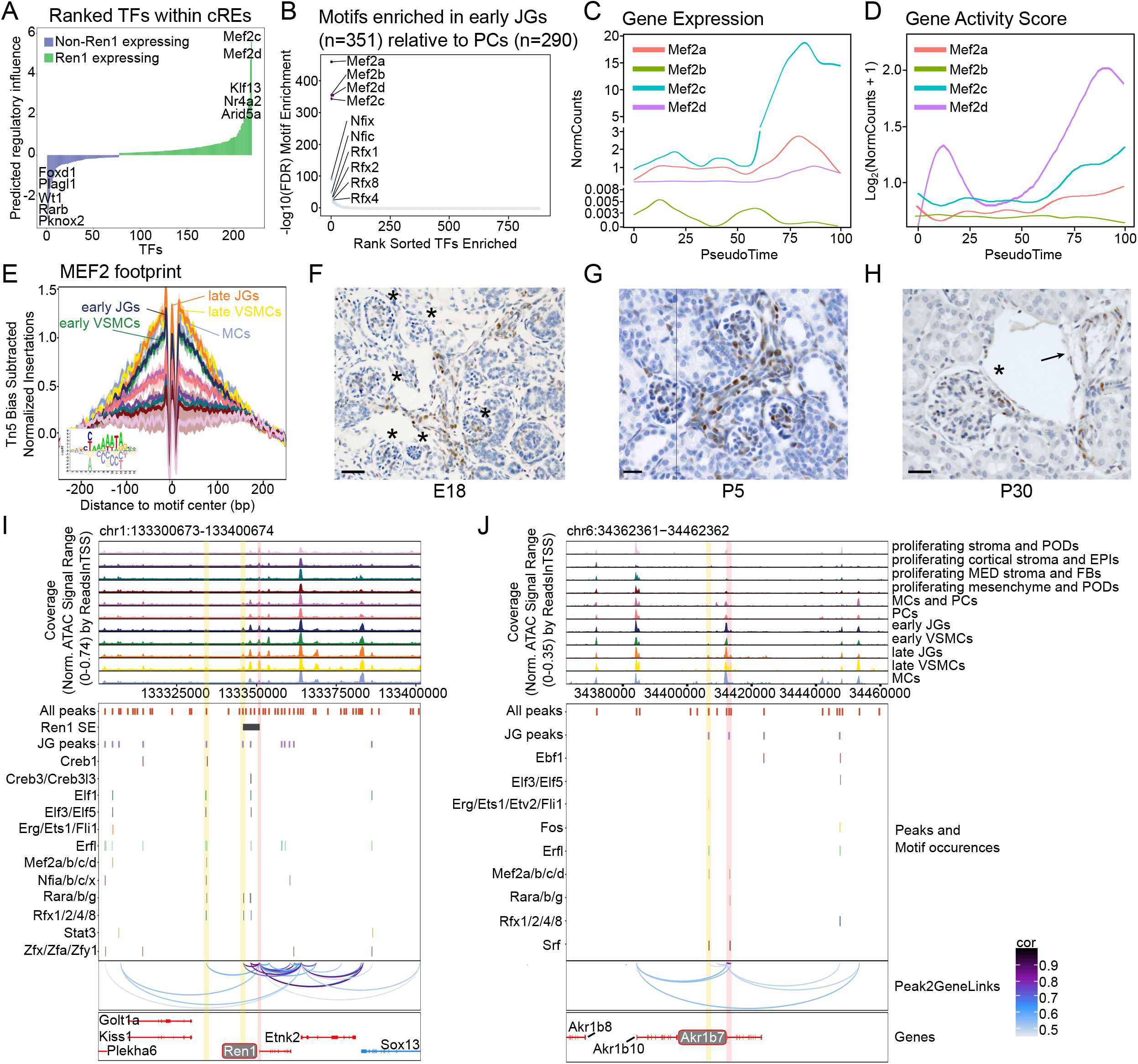
MEF2 family of TFs uniquely defines the JG population. (A) TFs ranked by predicted regulatory score related to Ren1 expression. (B) Top enriched motifs in early JG cells versus early PCs. MEF2 gene expression (C) or gene score activity (D) along pseudotime trajectory. (E) MEF2 family footprints are enriched in JG cells (Inset represents the motif logo). Immunohistochemistry for Mef2c in kidney sections at E18 (F), P5 (G), and P30 (H). The brown color stains Mef2c producing cells. In (F), asterisks highlight glomeruli with Mef2c along afferent arterioles, an interlobular arteriole, and at MCs. In (G), Mef2c is observed inside glomeruli, at MCs, afferent arterioles, an interlobular arteriole, and at the JGA. (H) At P30, the asterisk highlights a glomerulus with Mef2c in the MCs and JGA. A larger vessel (black arrow), depicts Mef2c within its walls. Scale bars: 50µm. Browser tracks identify preferentially enriched open chromatin in the JG cell clusters at (I) Ren1 and (J) Akr1b7. Motif occurrences of enriched TFs identify putative binding sites in open and co-accessible peaks. Yellow fill box highlights uniquely enriched peaks in JG clusters with co-accessibility to the Ren1 or Akr1b7 promoters. Pink fill box highlights core promoters. (cREs: cis-regulatory elements; EPIs: epithelial cells; FBs: fibroblasts; JGA: juxtaglomerular apparatus; JGs: juxtaglomerular cells; MCs: mesangial cells; MED: medullary; PCs: pericytes; PODs: podocytes; SE: super-enhancer; VSMCs: vascular smooth muscle cells)

With the data emphasizing a significant role for MEF2 family members in JG cell differentiation we investigated individual MEF2 members and found different patterns of gene activity scores and gene expression across differentiation. *Mef2a* peaked in late differentiation (Fig. 6C, D). Conversely, *Mef2b* peaked during the formation of early JG cells before fading during late JG maturation (Fig. 6C, D). Both *Mef2c* and *Mef2d* displayed overall similar patterns of activity, showing maximum accessibility and expression in the fully differentiated JG and VSMC populations although at different magnitudes (Fig. 6C, D). We also identified an overall enrichment of MEF2 TF footprints along the differentiation trajectory and in JG cells specifically (Fig. 6E).

To support the participation of MEF2 in renin cell development, we confirmed the presence of MEF2 members using immunohistochemistry (IHC) in one month old wild-type mice (*Mef2c* Fig. 6F-H) and using RNAscope in kidney sections of mice from E18, P5, and P30. IHC revealed the presence of *Mef2c* at the juxtaglomerular apparatus (JGA) and within VSMCs along vessels at the JGA (Fig. 6F-H). In addition, *Mef2b*, as revealed by RNAscope, is co-enriched in renin expressing cells throughout development (Fig. S2)

The participation of the MEF2 family in the muscular cells is easily recognized. However, some studies suggested that the MEF2 family are also transcriptional effectors of the vascular endothelial growth factor A (*Vegfa*) and Delta-like ligand 4 (*Dll4*)/Notch signaling with concomitant recruitment of the histone acetyltransferase (HAT) p300 for sprouting angiogenesis. The Notch signaling pathway and the recruitment of co-activators with HAT activity, such as the CBP/p300 complex, is essential for renin expression by JG cells and the conservation of a healthy kidney^29^. MEF2 target gene activation has been directly linked to stimulation by p300^74,75^, which is itself critical to remodeling of chromatin at the renin locus^7^. Clinically, *Mef2c* haploinsufficiency is often misdiagnosed as Rett syndrome. Indeed, there is overlap among *Mef2c* deficiencies between Rett syndrome, Angelman syndrome, Pitt-Hopkins syndrome, and CDKL5 deficiency disorder^76,77^. Although there is not a specific kidney phenotype, duplex kidney (or duplicated collecting system) is an anomaly that has been reported in one cohort of a pediatric population with *Mef2c* haploinsuficiency^77^. This highlights our findings of a novel function for *Mef2c* in normal kidney physiology.

We investigated whether any of the identified TFs have putative binding sites at *Ren1* or *Akr1b7* (Fig. 6I-J). Because open chromatin regions with shared accessibility may represent distinct regulatory networks, we identified co-accessible peaks genome-wide to provide a means to uncover features relevant to cell populations that make up the renin cell trajectory. This co-accessibility identifies chromatin regions with strong correlation across many cells and can indicate cell-type specific regulatory regions. We identified enriched motif binding sites present in regions unique to the JG cell populations at *Ren1*. Specifically, *Rfx1, Rfx2, Rfx4*, and *Rfx8* of the RFX family have motif occurrences present in peaks unique to JG cells and are enriched broadly in late differentiation clusters (Fig. S5E). The *Rfx2* motif has been previously identified in renin cell promoter and enhancer elements^14^. The RFX family regulates their target genes through a DNA sequence motif called X-box. They are usually involved in cellular differentiation and developmental processes^78^. The literature about the participation of this TF family in kidney physiology is scarce. Nevertheless, several human homologs of RFX target genes are known to be involved in diseases such as Bardet-Biedl syndrome^79^, whose kidney abnormalities are a major cause of morbidity and mortality. *Rfx2* influences HLA class II expression, but the importance of this to renin cells remains to be further investigated.

We also identified enriched motifs of zinc-family member proteins, *Zfa* and *Zfy1* (Fig. 6I; Fig. S5J). *Zfy1* (Zinc finger Y-chromosomal protein) is associated with kidney abnormalities including Frasier syndrome, which is characterized by progressive focal and segmental glomerulosclerosis, nephrotic syndrome, and kidney failure due to mutations of the Wilms tumor gene 1 (*Wt1*)^80^. The mechanism for JG cell differentiation requires further investigation.

We observed a number of ETS TF family members (*Elf1, Elf3, Elf5, Erg*, and *Erfl*) with significantly enriched motif occurrences present in regions that define the JG cell populations (Fig. 6I). Two of these members are known repressors: *Erfl* (ETS Repressor Factor-Like) and its paralog *Erf* (Ets2 Repressor Factor). We speculate these genes would orchestrate the ETS family activity via negative feedback. Some ETS family members are cell lineage specific, participating in cell development and differentiation by increasing related enhancer or promoter activities^81^. Usually, this family of TFs regulates target gene expression by interaction with additional TFs. The most described interaction is with Jun family proteins in the promoter of metalloproteinase (MMP) genes, with recruitment of CBP/p300 as co-activators.^81^. MMPs have been increasingly linked to both normal and abnormal pathology in the kidney, such as acute kidney injury, diabetic nephropathy, glomerular nephritis, inherited kidney disease, and chronic allograft nephropathy. They regulate extracellular matrix (ECM) degradation, but also inflammation, cell proliferation, angiogenesis, and apoptosis. They are typically expressed by resident macrophages or lymphocytes, but also by fibroblasts and MCs^82^. This is in accordance with previous studies that MCs are the major contributors to mesangial turnover. Dysfunctions in this process activate and release excessive TGF-*β*1, contributing to glomerular sclerosis and interstitial fibrosis^82^. Further, MCs play a role in glomerular contraction and filtration, thus affecting the glomerular filtration rate (GFR), although the molecular mechanisms regulating this process remain unclear^83–85^. Interestingly, some ECM components are linked to matrix-cell signaling in response to mechanical stretch, especially via *α*5*β*1 and *α*8*β*1 integrins. We described the participation of *β*1 integrins and lamin A/C in renin gene expression and renin secretion to maintain blood-pressure homeostasis, unravelling the long-sought renal baroceptor mechanism^86^. Potentially, a similar molecular mechanism would allow the MCs to activate genes related to a contractile phenotype (Fig. 7A) to enable them to alter the intraglomerular pressure and the ultrafiltration surface area, leading to a tight control of glomerular filtration rate (GFR).

The cAMP response elements, *Creb1, Creb3, Crem*, and *Creb3l3*, are also enriched in JG marker regions (Fig. 6I; Fig. S5G). *Creb* is known to switch “on” the renin gene^87^. *Creb3* is also present in later stages of JG cells differentiation, which is in full agreement with previous studies. *Creb* is itself activated by the PKA (Protein kinase A)/cAMP pathway, one of the major regulators of renin synthesis and release^6,12,22,23,87^.

We observed significant motif occurrences for MEF2 members in a chromatin region unique to the JG cell populations with co-accessibility to the *Ren1* core promoter. (Fig. 6I; Fig. S5A). We found further support of the contribution of MEF2 members by observing enrichment within open chromatin of JG populations at the *Akr1b7* locus (Fig. 6J). The RAR, RFX, and ETS families are also enriched at *Akr1b7* as at *Ren1*, supporting these TFs as being important contributors to renin cell formation and identity. With *Rarb* predicted to be inversely related to renin activity (Fig. 6A), we would predict that RAR members downregulate *Ren1* and *Akr1b7* expression via binding sites identified at both loci (Fig. 6I-J). *Srf*, which displays an enriched footprint in JG cell populations (Fig. 5C), has binding sites at the *Akr1b7* promoter and within co-accessible regions (Fig. 6J).

### JGs and VSMCs share a dual endocrine and contractile phenotype

Past work has highlighted the bivalent nature of renin expressing JG cells between endocrine and contractile phenotypes^14^, and here again we emphasize the presence of markers of both phenotypes present in JGs and VSMCs (Fig. 7A). The consistent enrichment of the MEF2 family across developmental timepoints suggest they are the differentiation drivers for the *FoxD1* descendants’ cells. *Mef2c* has an essential role for skeletal muscle growth and differentiation, and we observed significant increases in transcript abundance and accessibility for *Mef2c* by the formation of early JGs (Fig. 7A). Furthermore, we observed enrichment of *Srf* in P30 populations of JGs and VSMCs. *Srf*, a paralog of *Mef2c*^88^, has been shown to be involved in VSMC development and maintenance and likely contributes to the plasticity of cells able to express renin^14^ (Fig. 7A). Strikingly, motif enrichment in VSMCs with and without renin expressing cells was very similar and the overlap in accessibility and expression patterns between these cell populations is strong (Fig. 7A). Both showed the MEF2 family as the highest enriched motifs. Spatial transcriptomics with precise refinement and resolution would provide additional aid to resolve these similarities in future work.

**Fig. 7:**
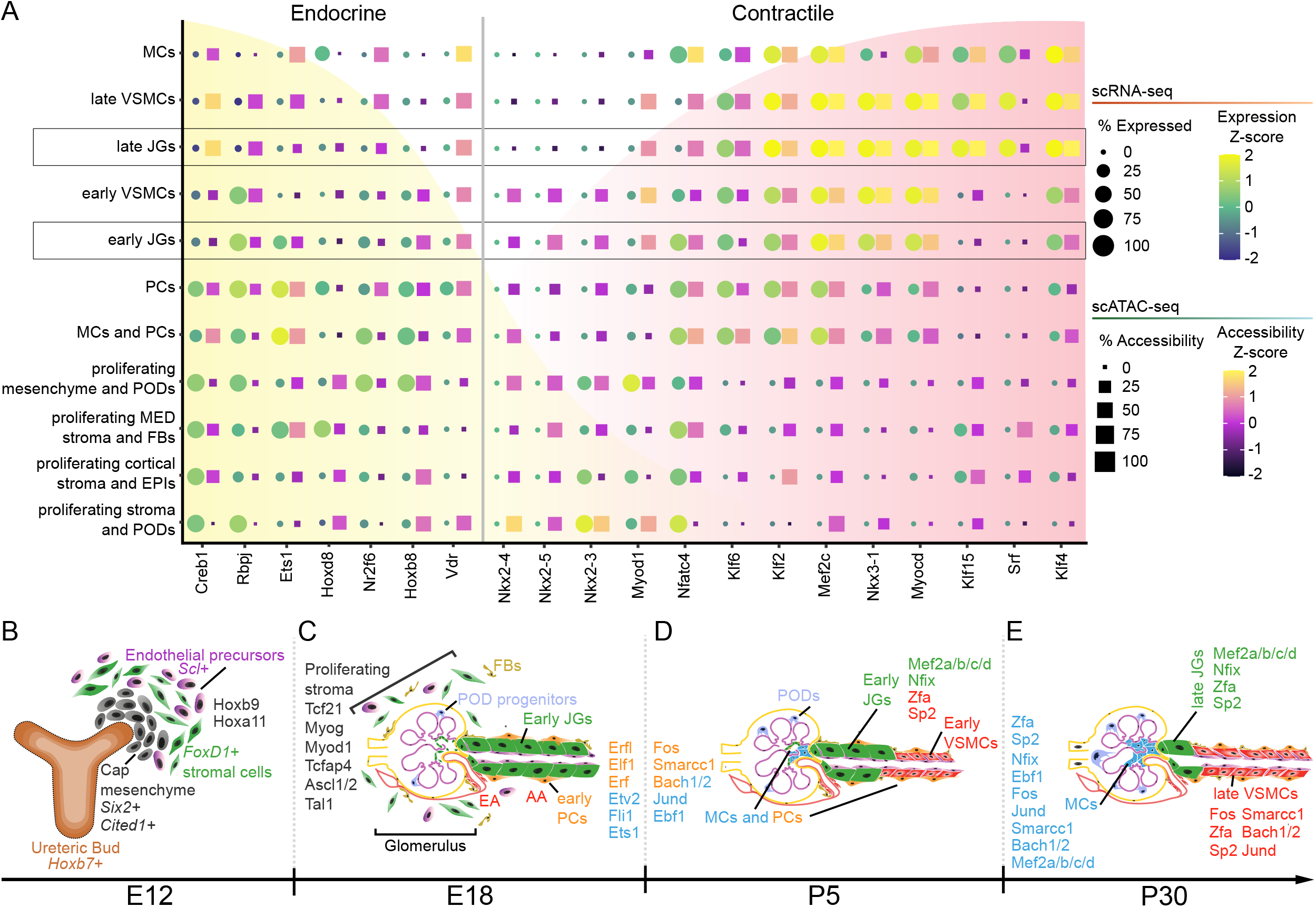
Trajectory analysis reveals complexity of JG cell phenotype and differentiation. (A) Dot plot of gene activity scores and gene expression values for markers of endocrine or contractile phenotypes. JG cell differentiation trajectory highlights enriched motifs (from Fig. 3B) in each cell population (B-E). (B) Hoxb9 and Hoxa11 are enriched in the E12 populations of proliferating stroma and precursor cell populations. (C) Tcf21, Myog, Myod1, Tcfap4, Ascl1 and Ascl2, and Tal1 are enriched in the E18 proliferating stroma populations. In the early PCs and mixed MCs and PCs populations present in the transition between E18 to P5, Erfl, Elf1, Erf, Etv2, Fli1, Fos, Smarcc1, Bach1 and Bach2, Jund, and Ebf1 are significantly enriched. (D) In the early JGs and early VSMCs there is significant enrichment of Mef2a/b/c/d, Nfix, Zfa, and Sp2 motifs. (E) By P30, the MCs are enriched for Fos, Smarcc1, Bach1 and Bach2, Jund, and Ebf1, and to a lesser extent Mef2a/b/c/d, Nfix, Zfa, and Sp2. The late JGs are enriched for Mef2a/b/c/d, Nfix, Zfa, and Sp2 motifs. In contrast, the late VSMCs are enriched for Fos, Smarcc1, Bach1 and Bach2, Jund, Ebf1, Zfa, and Sp2. (AA: afferent arteriole; EA: efferent arteriole; EPIs: epithelial cells; FBs: fibroblasts; JGs: juxtaglomerular cells; MCs: mesangial cells; MED: medullary; PCs: pericytes; PODs: podocytes; VSMCs: vascular smooth muscle cells)

## Discussion

Nephrogenesis is a complex process, and in mice starts around embryonic day E11.5 and finishes around 3-5 days after birth^89^. The vertebrate kidney derives from the intermediate mesoderm, which differentiates into three successive embryonic kidney layers: the pronephros, the mesonephros, and the metanephros. The metanephros develops into the permanent, mature kidney in mammals, and results from the reciprocal interaction between the ureteric bud and the metanephric mesenchyme (Fig. 7B). The former will generate the tubular system and glomerular epithelia^89^. The latter can be divided into condensing and loose mesenchyme. The loose mesenchyme cells express the TF, *FoxD1*, and are the precursors of all the renal VSMCs, renin expressing JG cells, MCs, and renal interstitial PCs (Fig. 7B-E). The loose mesenchyme also contains endothelial precursors, typically marked by the expression of *Scl* (Stem-Cell Leukemia), which may themselves transiently express *FoxD1* (Fig. 7B-E). Thus, the presence of endothelial cells in our observed cell populations is not unexpected. Interestingly, renin precursors *per se* give rise to VSMCs, MCs, and PCs capable of dedifferentiating and re-expressing the renin phenotype under homeostatic challenge^90^.

We employed high-throughput 10X-based scRNA-seq and scATAC-seq technology to simultaneously measure the transcriptome and accessibilome of progenitors and mature renin expressing cells in the mouse kidney. In this study, we uncovered how successive changes in chromatin states -opening and closing of specific domainsand sets of TF combinations along the genome instruct the *FoxD1* progenitor’ cells to adopt the different cell fates that populate and assemble the kidney vasculature. Fig. 7B-E illustrates the point with respect to the TFs that characterize each cell state of differentiation as the kidney arterioles mature and the cells that compose them move along their orchestrated differentiation pathway. For instance, proliferating stroma utilizes a different set of TFs from the early JG cell, the late VSMCs, or the MCs prior for which there were practically no unique or sets of unique TFs that identified them as such. To achieve this in-depth granular understanding required a deeper bioinformatic analysis that dealt with the question of what constitutes cell “identity.” This was made even more difficult as the cells were undergoing transitional and ever-changing states of maturation/differentiation.

To reveal a nuanced understanding of cell fate decisions, we could not rely exclusively on unsupervised cell clustering techniques. As a consequence of shared lineage and the renin phenotype switching role of cells of this background, identifying JG cells required the manual annotation of JG populations. We did this first by identifying cells with renin expression in the 90th percentile and with gene activity scores in the upper quartile. We further subdivided that population by separating early forming JG cells, as those cells falling in the aforementioned percentiles that formed prior to P30. Because our goal was to understand differences along differentiation, this enabled us to identify changes between cell populations both broadly and at specific transition points during development. We similarly identified unique populations of VSMCs by leveraging canonical VSMC markers and annotating cells with combined expression and gene activity scores of those markers in the upper quartiles, and split early versus late forming VSMCs in the same way as the JG cell population. Thus, we achieved the necessary granularity to unravel differences between these cells of a shared lineage and potential shared functionalities. A similar strategy in future studies of rare or phenotypically similar cell populations will be essential.

The results of this study have additional medical implications: understanding the regulatory landscape of renin expressing JG cells is necessary to better learn the control and function of this rare cell population as overactivation or aberrant activity of the RAS is a key factor in cardiovascular and kidney pathologies^5,6,22,23,91–96^. Our current understanding of the epigenomic regulatory landscape in renin expressing JG cells is limited, therefore it is urgent we expand our knowledge to understand consequences of disruptions to the RAS that occur in human health and disease and the effects of drugs targeting this pathway, commonly used to treat hypertension.

In summary, we have provided the first developmental trajectory of renin cell differentiation as they become JG cells in a single-cell atlas of kidney vascular open chromatin and highlighted novel factors important for their stage-specific differentiation during embryonic and postnatal life (Fig. 7B-E). Specifically, the MEF2 family of transcription factors were enriched in late differentiation clusters, including the JG cell populations (Fig. 7D, E). Previous studies of the importance of MEF2 members in angiogenesis^75^, and the interaction of MEF2 members with p300 which we identified as crucial for renin cell identity^97,98^, provide a foundation for future *in vivo* and *in vitro* experiments to directly interrogate the individual roles of MEF2 members and other stage-specific TFs described herein. The data generated by the present atlas would be useful to explore whether “waves” of TFs coordinate their action(s) to guide renin cells through specific developmental stages as they form the classical JG cell.

## Article Information

## Non-standard Abbreviations and Acronyms

AA: afferent arteriole
CD: collecting duct
E12: embryonic day 12
E18: embryonic day 18
EA: efferent arteriole
ENDOs: endothelial cells EPIs epithelial cells
FBs: fibroblasts
GFR: glomerular filtration rate
ITSs: interstitial cells
JGA: juxtaglomerular apparatus
JGs: juxtaglomerular cells
MCs: mesangial cells
MED: medullary
MEF: myocyte enhancer factor
NFIX: Nuclear Factor I X
P5: five days old
P30: one month old
PCs: pericytes
PCTs: proximal convoluted tubules
PODs: podocytes
SE: super-enhancer
VSMCs: vascular smooth muscle cells

## Authors’ Contributions

RAG and MLSSL conceived and designed the project. AGM designed and performed animal experiments and library preparation. JPS and AGM discussed and performed computational data analysis. AGM analyzed single time points and JPS developed methods to identify JG cells and evaluate differentiation trajectories across all developmental timepoints. NCS provided bioinformatic support. SM made libraries and deleted the *Nfix* gene in As4.1 cells. JPS wrote the manuscript and all authors discussed the results and contributed to the final manuscript.

## Acknowledgments

The authors gratefully acknowledge the technical and analytical help of Minghong Li, Xiuyin Liang, Fang Xu, Devon Farrar and Thomas Wagamon. We thank Vidya Nagalakshmi for preliminary protocols of nuclei isolation.

## Supplemental Material

- Supplemental Methods.
- Figures S1-S7
- References^99–125^

## Materials and Methods

### Mouse models

All animals were maintained in a room with controlled temperature and humidity under a 12-hour light/dark cycle. All animals were handled in accordance with the National Institutes of Health guidelines for the care and use of experimental animals, and the study was approved by the Institutional Animal Care and Use Committee of the University of Virginia.

To generate *FoxD1cre/+*;*R26R-mTmG* mice, we crossed *FoxD1cre/+* mice with the *R26R-mTmG* mice^40,99^. scRNA-seq and scATAC-Seq were performed using the 10X Genomics technology according to the respective protocol^100^. For the scATAC-Seq, we performed the nuclei isolation following the manufacturer protocol^101^, and all the experiments targeted at least 2000 nuclei, while the scRNA-seq always targeted more than 1000 cells. The experiments were performed at specific time points of the mouse kidney development: E12 (embryonic day 12), E18 (embryonic day 18), P5 (five days old) and P30 (one month old). Please, note that mice nephrogenesis and vascular development starts at E11.5 and continues after birth for about 3-7 days, respectively^89^.

### Isolation of kidney cells

#### Isolation of kidney single cells: E12

Pregnant mice were injected with tribromoethanol at enough dose to keep alive but anesthetized. Then, the small fetuses were removed one by one. A small squash from the lung was used to identify the GFP pups, using an EvosFLC cell imaging system (LifeTechnologiesTM, California, USA). Once they were identified, the metanephros area was removed under microscopy and harvested in ice cold dPBS. The tissue was then minced carefully with a razor blade and transferred to 1.7mL Eppendorf tube. 300 *µ*L of TryPLETM Express (Gibco, New York, USA) was added and incubated at 37^◦^ for 5 minutes. Then, 600 *µ*L of DMEM + 5% FBS was added to the Eppendorf tube. The mixture was homogenized by pipetting up and down. The solution was then filtered with a 40 *µ*m nylon cell strainer. Tissue chunks were removed from the mesh surface, placed again in an Eppendorf tube and the process repeated a second time. Meanwhile, the flow-through filtrate was placed in a new Eppendorf tube and centrifuged at 4^◦^C, 150g for 5 minutes. The supernatant was removed and the pellet resuspended with resuspension buffer. All the tubes were combined in the end and the cells were ready for single-cell capture.

#### Isolation of kidney single cells: E18, P5 and P30

Animals were anesthetized with tribromoethanol (300 mg/kg). P5 and P30 mice kidneys were excised and decapsulated. Then, the kidney cortices were dissected, minced with a razor blade, and transferred into a 15 mL tube with 5 mL of enzymatic solution (0.3% collagenase A [Millipore-Sigma], 0.25% trypsin [Millipore-Sigma], and 0.0021% DNase I [Millipore-Sigma]). The tubes were placed inside a shaking incubator (80 RPM) for 15 minutes at 37^◦^C. The solution was pipetted up/down 10 times with a sterile transfer pipette and allowed to settle for 2 minutes, and the supernatant was collected in a fresh tube on ice. The enzymatic solution was added to the 15 mL tube containing the remaining undigested cortices, and the digestion procedure was repeated a total of 3 times. The supernatants were pooled and centrifuged at 800 g for 4 minutes at 4^◦^C using a Sorvall RT7 refrigerated centrifuge (Sorvall, Newtown, CT). The cell pellet was resuspended in fresh buffer 1 (130 mM NaCl, 5 mM KCl, 2 mM CaCl2, 10 mM glucose, 20 mM sucrose, 10 mM HEPES, pH 7.4), and the suspension was poured through a sterile 100 *µ*m nylon cell strainer (Corning Inc., Corning, NY) and washed with buffer 1. The flow-through was poured through a sterile 40 *µ*m nylon cell strainer (Corning Inc.) and washed with buffer 1. The flow-through was centrifuged at 1,100 g for 4 min at 4^◦^C. The cell pellet was resuspended in 1.5 mL of resuspension buffer [PBS, 1% FBS, 1 mM EDTA, DNAase I (Millipore-Sigma)]. DAPI (Millipore-Sigma) was added to the cells to identify the living cells. The GFP positive cells were collected by Fluorescent-Activated Cell Sorting (FACS)^6^ and resuspended in DMEM (Dulbecco’s Modified Eagle Medium, Gibco, Netherlands) with 10% FBS (Fetal Bovine Serum) for immediate use. The sorters were either an Influx Cell Sorter (Becton Dickinson, Franklin Lakes, NJ) or a FACS Aria Fusion Cell Sorter (Becton Dickinson), both located at the Flow Cytometry Core Facility at the University of Virginia.

### scATAC-seq library preparation

The initial steps of capture between scATAC-seq and scRNA-seq are similar. The GFP-positive cells were washed in dPBS with 0.04% BSA twice, and the sorted GFP-positive cells were washed in dPBS with 0.04% BSA twice. The cells were counted with the CellometerMini (Nexcelom Bioscience, Massachussets, USA). Sorted cells with viability higher than 80% and absence of clumps were chosen to proceed. However, the scATAC-Seq protocol requires nuclei isolation. Our experiments yielded less than 100,000 cells. Therefore, we have used the protocol designed for Low Cell Input Nuclei Isolation. Briefly, we centrifuged the cells at 500g at 4^◦^C for 10 minutes, removed the supernatant carefully and resuspended the cells in a freshly prepared lysis buffer (1M Tris-HCl, 5M NaCl, 1M MgCl2, 10% BSA, 10% Tween-20). We have optimized the lysis incubation time for 7 minutes on ice. Next, we immediately added freshly made washing buffer (1M Tris-HCl, 5M NaCl, 1M MgCl2, 10% Tween-20, 10% Nonidet P40, 5% digitonin and 10% BSA). This washing was performed twice, and then supernatant was removed for a final wash with diluted nuclei buffer (nuclei buffer 20X, 10X Genomics, diluted in nuclease free water to 1X). Once this was completed, we again removed the supernatant, resuspended the nuclei pellet and counted them with a Neubauer Chamber (Spencer, Buffalo, USA). In all our experiments we were able to target between 1000-5000 nuclei. Nuclei were loaded and captured using the Chromium System (10X Genomics, Pleasanton, CA) following the manufacturer’s recommendation^100^ using the Chromium Next GEM (Cell-Gel Beads in Emulsion) Chip H with reagents of Chromium Next GEM Reagent Kits v1.1 (10X Genomics). Initially, nuclei were incubated with the transposition mix, that includes a transposase, and later the GEMs were barcoded and PCR amplified to generate the cDNA. The cDNA was then cleaned with Dynabeads and SPRIselect, end-repaired, adaptor-ligated and amplificated by PCR. The constructed libraries were sequenced on an Illumina HiSeq 2500/4000 platform (150-bp paired-end reads).

### scRNA-seq library preparation

Sorted GFP-positive cells were spun down at 500g for 10 minutes in a Sorvall RT7 refrigerated 4^◦^C centrifuge (Sorvall, Newtown, CT). Then, the supernatant was carefully removed, and the cell pellet was resuspended in dPBS (Dulbecco’s Phosphate Buffered Saline, Gibco, UK) with 0.04% BSA (UltraPureTM Bovine Serum Albumin, Invitrogen, Lithuania) 10 times with wide-bore tips. The process was repeated once, and the cells were counted with the CellometerMini (Nexcelon Bioscience, Massachussets, USA). Experiments with more than 500 cells/*µ*L, viability higher than 85%, and absence of clumps were chosen to proceed. Single cells were loaded and captured with the Chromium System (10X Genomics, Pleasanton, CA) following the manufacturer’s recommendation^100^ using the Chromium Next GEM Chip G with reagents of Chromium Next GEM Single Cell 3^′^ Reagent Kits v3.1 (10X Genomics). The GEMs were generated and incubated to generate the barcoded cDNA. cDNA was cleaned with Dynabeads and amplified by PCR. cDNA was then enzymatically fragmented, end-repaired, poly-A tailed, adaptorligated, and amplificated by PCR. The constructed libraries were sequenced on an Illumina HiSeq 2500/4000 platform (150-bp paired-end reads).

### Read preprocessing

#### Genome and transcriptome annotations

We performed all downstream analyses using the mm10 genome. For Cell Ranger pipeline analyses, we used the `refdata-cellranger-arc-mm10-2020-A-2.0.0` reference data and for R-based analysis, the BSgenome package `BSgenome.Mmusculus.UCSC.mm10` was used.

#### scATAC-seq alignment and fragment matrix generation

We processed fastq files from the 10X Genomics Single Cell ATAC platform using the Cell Ranger ATAC pipeline (version: `cellranger-atac-2.0.0)`. Cell Ranger trims primer sequences using a modified cutadapt tool^102^ before alignment using a modified bwa-mem algorithm^103,104^. Duplicate reads were removed based on the start, end, and hashed barcode of aligned reads. Finally, read fragments were corrected for Tn5 transposase binding biases and stored as position-sorted fragment files.

#### scATAC-seq quality control

We processed Cell Ranger fragment files using the `ArchR v1.0.1 R` package^105^. Arrow files were produced from each time point’s fragments file. We determined the presence of putative cell doublets using the ArchR function addDoubletScores, which determines doublet identity by embedding synthetically created pseudo-doublets from the input data into the shared sample space. We removed cells that behaved like pseudo-doublets (n=599) prior to downstream analysis (Figure S7A-D). We further excluded cells with TSS enrichment scores less than 10 as these low TSS scores are indicative of low signal-to-noise and poor quality (Figure S7E-H)^126^. Finally, we evaluated fragment distributions for each developmental time point to confirm the expected periodicity of nucleosomal positioning (Figure S7I-L).

#### scATAC-seq dimensionality reduction and batch correction

Following quality control filtering, an iterative Latent Semantic Indexing dimensionality reduction^106,107^ technique using Singular Value Decomposition (SVD) to embed the most valuable sample information in low dimensional space^105^ was applied (see ArchR `addIterativeLSI`). This reduced dimensionality matrix was further corrected for batch effects using the Harmony^108^ algorithm originally developed for scRNA-seq data but extended to scATAC-seq in ArchR (using the function addHarmony)^105^.

#### scATAC-seq cell clustering

We then clustered the batch-corrected reduced dimensionality matrix to identify cells based on shared chromatin accessibility patterns using the `addClusters` function in ArchR^105^. These cluster embeddings were visualized using Uniform Manifold Approximation and Projection (UMAP) to evaluate dimensionality reduction and batch correction results across the integrated time points.

Because scATAC-seq data is inherently sparse, pseudo-bulk replicates were generated (`addGroupCoverages` function) by grouping similar single cells into aggregate profiles similar to bulk ATAC-seq data and then calling peaks. Peak calls were generated using MACS2^109^ with the addReproduciblePeakSet function in ArchR.

#### scRNA-seq alignment and feature-barcode matrix generation

Next, we processed scRNA-seq fastq files from the 10X Genomics Single Cell Gene Expression platform using the Cell Ranger pipeline (version: `cellranger-6.0.1`). The Cell Ranger pipeline trims non-template sequence and aligns reads using the STAR aligner to genomic and transcriptomic annotations^127^. Confidently mapped transcriptomic reads are grouped by barcode, UMI, and gene. The number of reads mapped to each gene was calculated using UMI-based counts. Finally, filtered UMIs were mapped to barcodes to build feature-barcode matrices for downstream analysis.

#### scRNA-seq quality control

We processed the filtered feature-barcode matrices produced by Cell Ranger in R using `Seurat`^110^. Abnormally low (*<*200) or high (E12 *>*9000; E18*>*7000; P5*>*7500; P30*>*4500) numbers of features (Figure S7M-P) indicative of low quality and low information or doublet cells respectively were removed^111,112^. While a stringent threshold (*<*5%^128^) for mitochondrial contamination is often recommended, cells with high energy demands may contain higher than expected mitochondrial sequence^113,114^. Based on the experimental design of capturing actively dividing kidney cells, we loosened this threshold to remove cells with *>*30% mitochondrial reads (Figure S7Q-T) to retain high quality samples. Finally, hemoglobin mapped reads indicative of red blood cell contamination are restricted to retain cells with *<*30% hemoglobin (Figure S7U-X).

#### scRNA-seq cell clustering

Next, scRNA-seq cells were normalized (see Seurat `NormalizeData`) by log-transforming the scaled (number of features divided by library size and multiplied by a scaling factor of 10000) read counts + 1^110,115^. We merged each scRNA-seq time point and identified the top 2000 most variable features (see Seurat `FindVariableFeatures`) between cells. We applied a linear transformation (see Seurat `ScaleData`) to scale expression to a mean of 0 between cells with a maximum variance of 1 to reduce the dominance of highly-expressed genes prior to dimensionality reduction.

Next, we performed Principal Component Analysis (PCA) on the normalized data using the previously identified top features (see Seurat `RunPCA`). To reduce technical noise, we performed a JackStraw^129^ procedure (see Seurat JackStraw) which randomly subsamples the data, re-performs PCA, and identifies the components with the highest enrichment of low p-value features. We then implemented a graph-based clustering approach to identify groups of similar cells which we subsequently visualized using UMAP (see Seurat `RunUMAP`).

#### scRNA-seq cell identification

We annotated individual cell clusters using marker genes defining each cluster. Specifically, the top 25 marker genes for each cluster were identified by finding differentially expressed genes being present in at least 25% of the cells between groups with a 0.25 log fold change between clusters for that gene (see Seurat `FindAllMarkers`). These uniquely expressed marker genes were used to annotate individual clusters. To perform this annotation, an iterative approach was performed by comparing the top markers to previously identified markers from a series of mouse cell atlases and a commercial database of cell markers (cellKb^116^). First, internally identified markers from internal data was overlapped with top markers from identified clusters. Second, markers were overlapped with cell identity markers from three atlases of developing mouse kidney cells^117–119^. Third, top identified markers were matched with cell signatures using the commercial CellKb database to create rank ordered lists of top matching gene signatures to cell signatures. The resulting top ranked hits from all three methods were then merged and cell cluster identities manually annotated.

### Integrating transcriptome and accessibilome

We performed label-transfer to integrate scRNA-seq annotated data with the scATAC-seq data. Anchors between gene scores, a measure of gene expression based on chromatin accessibility, and RNA expression from the scRNA-seq data was combined through the `addGeneIntegrationMatrix` function from `ArchR`. First, a gene score matrix is generated by summing reads in peaks with the nearest annotated gene to calculate gene activity by proxy of chromatin accessibility^105^. With this information, cells with the most similar gene score derived expression profile are matched to scRNA-seq expression. This has the intended benefit of labeling ATAC-derived clusters with the RNA-seq based cell annotations. With this integration, we investigated the chromatin accessibility of individual cell types to identify cell-type specific and co-accessible peaks, differentially accessible regions, enriched motifs, and cellular trajectories.

### Renin cell differentiation trajectory

To identify a renin cell differentiation traecjtory, we performed trajectory inference to uncover the dynamic chromatin and expression changes that lead from *FoxD1*+ progenitor cells to renin expressing cells during mouse kidney development. We calculated a pseudotime trajectory for each cell. Cells early in the renin cell differentiation trajectory were identified by filtering cells on high *FoxD1*+ expression from scRNA-seq and gene score activity from scATAC-seq data. We then identified cells enriched (both integrated RNA-seq expression and gene score activity) for the renin marker genes, *Ren1* and *Akr1b7*. Clusters that fulfilled these requirements were utilized as a backbone of ordered vectors and a pseudotime trajectory constructed using ArchR’s addTrajectory function. This trajectory, and relevant markers, were then visualized on the UMAP embeddings of the LSI and Harmony corrected reduced matrix using the function `plotTrajectory`.

#### Identifying renin cells

To determine the identity of the renin expressing cells, we evaluated markers of mature renin cells and VSMCs. Both renin cells and VSCMs are derived from a shared lineage (i.e. *FoxD1* progenitors). To isolate mature renin cells, our representative JG populations, we exclude cells with predominant VSMC markers. Using MAGIC^120^ imputed gene scores and gene expression values from the scATAC-seq and scRNA-seq respectively, we isolate individual cells with gene expression values greater than the 90th percentile and gene score values greater than the 75th percentile for the JG markers, *Ren1* and *Ark1b7*. For VSMCs, we perform the same filtering based on the VSMC markers *Acta2, Myh11*, and *Cnn1* but including the upper quartile of both gene expression and gene score values. We then take the difference between the identified JG cells and VSMCs to create renin cell exclusive subpopulations, with the remainder representing mature VSMCs. We further subdivide these populations based on the time of formation during development. Populations forming prior to P30 are labeled “early JGs” or “early VSMCs” respectively, and the P30 populations labeled as “late JGs” or “late VSMCs”.

#### Renin trajectory marker peaks

We subsetted the clusters comprising the renin cell trajectory and identified the regulatory regions that differentially identify each subpopulation by adding pseudo-bulk replicates that recapitulate the biological variability within each cluster (`addGroupCoverages`) prior to calling peaks (`addReproduciblePeakSet`). To identify features that were differentially expressed or accessible between clusters, the `getMarkerFeatures` (ArchR) function was used and features with a false discovery rate of less than 0.1 and absolute log2 fold changes greater than 1 were identified. We then plotted marker genes based on integrated scRNA-seq and scATAC-seq data on the UMAP embedded reduced dimensionality matrix.

#### Motif annotations and enrichment

We added motif annotations in enriched chromatin regions using the cisbp^121^ database of annotated motifs through the `addMotifAnnotations` function of ArchR. This method utilizes chromVAR^122^ to calculate a TF deviation zscore that indicates relative enrichments of motifs in peak regions compared to background^105,122^. We leverage these deviation scores to uncover differentially accessible regions and motifs between cell clusters of the renin cell trajectory.

#### TF footprinting

To perform footprinting, motif positions were extracted from the renin cell trajectory clusters using `getPositions` (ArchR). These footprints were calculated using pseudo-bulk replicates of scATAC-seq data along the trajectory and normalized by subtracting the Tn5 insertion bias from the footprinting signal. Footprints of enriched motifs in each renin cell trajectory cluster were then plotted using `plotFootprints` (ArchR).

#### Regulatory network inference

Cell clusters were split into two broad populations based on *Ren1* expression and activity. We then employed the R package [RENIN](https://github.com/nledru/RENIN) to predict the regulators of *Ren1* differential expression. We included all predicted cis-regulatory elements to identify potential regulatory TFs. We next identified enriched motifs within the predicted cis-regulatory elements and ran an adaptive elastic net to predict which of these TFs likely regulate *Ren1*. Finally, we ranked TFs by their predicted regulatory score which `RENIN` calculates as the sum of estimated regulatory coefficients across all modeled genes that is weighted by the individual expression of those genes.

#### Peak co-accessibility and peak to gene links

We calculated open chromatin region co-accessibility using the `addCoAccessibility` function in ArchR. With integrated scRNA-seq data, we additionally looked for correlations between both accessibility and gene expression profiles between marker peaks using `getPeak2GeneLinks` at a resolution of 10000 to reduce the total number of links to prevent over-fitting^105^. These peak-gene linkages include not only correlated peaks but genes whose expression is also correlated between cells. The resulting co-accessible peaks and genes were visualized on browser tracks using the plotBrowserTrack function. To visualize how accessible regions and gene expression change along the trajectory, we also plotted a heatmap of these linked regions and genes using the `plotPeak2GeneHeatmap` function (ArchR).

We extended visualization in browser tracks by also plotting putative TF binding sites in regions of interest by looking for motif occurrence enriched above random using FIMO^123^. TF binding profile position frequency matrices (PFM) for motifs enriched in marker peaks were obtained from the exported cisbp database used within ArchR. For a wide region around the *Ren1* gene, enriched motif PFMs were used to find individual motif occurrences in this sequence and plotted alongside tracks of chromatin accessibility and co-accessible peaks. These binding sites were loaded into R as individual GRanges objects^124^ and concatenated into a single feature object and visualized in the browser by leveraging the features parameter of ArchR’s `plotBrowserTrack`.

#### Positive transcription factor regulators

Since the specific DNA motifs between related TFs are similar, linking individual TF gene expression with motif enrichments enables the identification of positively correlated genes with their matched motif. This was performed by first calculating the most variable TF deviation z-scores between clusters and then correlating those z-scores with the integrated gene expression from the scRNA-seq data `(correlateMatrices)`. Positive TF regulators were considered as those TFs with motif and gene expression correlation greater than 0.5, p-values less than 0.01, and whose maximum difference in z-score between clusters is in the first quartile. We also identified positive TF regulators derived from the gene score activity from scATAC-seq to identify positively correlated TFs based on motif enrichment and gene score `(correlateMatrices)`.

### RNAscope Assay

RNAscope was performed using the manufacturer protocol Multiplex Fluorescence v2 kit, ACDBio (Oxford, United Kingdom). ACDBio designed the probes, which were hybridized in a HybEz oven at 40^◦^C with 5 *µ*m thickness kidney sections which were formalin fixed and paraffin embedded. Following hybridization, signal development (HRP channel binding) and fluorescence was achieved using TSA plus© fluorescein (green) and TSA plus© Cyanine 5 (far red) fluorophores (PerkinElmer). Sample slides were treated with the nuclear stain DAPI and mounted with Prolong Gold Antifade mounting media.

### Immunohistochemistry

Formalin fixed and paraffin embedded kidneys sections were deparaffinized, rehydrated, and treated with 0.3% hydrogen peroxide in methanol. After blocking with 3% BSA and 2% horse serum, sections were incubated with anti-*Mef2c* antibody (rabbit monoclonal antibody) at 4^◦^C overnight. After washing, sections were incubated with biotinylated, secondary, goat anti–rabbit IgG antibody at room temperature for 30 minutes. Staining was amplified using the Vectastain ABC kit (Vector Laboratories) and developed with 3,3^′^-diaminobenzidine (MilliporeSigma). The sections were counterstained with hematoxylin (MilliporeSigma), dehydrated, and mounted with Cytoseal XYL (Thermo Fisher Scientific).

### Microscopy

Kidney sections were visualized using a Zeiss Imager M2 microscope equipped with an ApoTome-2 fitted with the AxioCam 305 color and AxioCam 506 mono camera (Zeiss).

### Cell culture

For CRISPR-Cas9 experiments, we used the renin-expressing As4.1 cell line (ATCC, CRL-2193)^125^. Cells were maintained in high glucose DMEM supplemented with 10% fetal bovine serum (both from Thermo Fisher Scientific) at 37^◦^C, 5% *CO*_2_.

#### CRISPR gene knockout

We performed knockout of *Nfix* in As4.1 cells using CRISPR-Cas9 ribonucleoproteins (RNPs). sgRNAs were provided with the Synthego Gene Knockout kit v2 (Synthego, Redwood City, CA) and Alt-R© S.p. HiFi Cas9 Nuclease V3 was from IDT (Integrated DNA Technologies, Coralville, IA). The Cas9 protein and sgRNAs (240 *ρ*mol) were mixed and incubated at room temperature for 20 minutes to generate RNPs. One million cells were resuspended in 100 *µ*L of Nucleofector solution SE (SE Cell Line 4D-NucleofectorTM X Kit (Lonza, Basel, Switzerland) and mixed with RNPs and Alt-R© Cas9 Electroporation Enhancer (IDT). As a negative control, cells were treated in the same way, except RNPs were excluded from the mix. To estimate the transfection efficienty, we nucleofected cells mixed with pmaxGFP plasmid. Cells were electroporated using the 4D-Nucleofector system (Lonza) under program CMAfter nucleofection, the cells were cultured in growth medium, passaged once, and analyzed 6 days after nucleofection.

#### RNA extraction and quantitative RT-PCR

We extracted total RNA from cultured cells using TRIzol reagent (Thermo Fisher Scientific) and the RNeasy Mini Kit (Qiagen, Dusseldorf, Germany). Reverse transcription (RT) was performed using oligo(dT) primers and M-MLV Reverse Transcriptase (Promega, Madison, WI) at 42^◦^C for 1 hour according to the manufacturer’s instructions. Quantitative real-time PCR was conducted using SYBR Green I (Thermo Fisher Scientific) and a CFX Connect system (Bio-Rad Laboratories, Hercules, CA). The forward primer for *Ren1* was 5’-ACAGTATCCCAACAGGAGAGACAAG-3’, and the reverse primer was 5’-GCACCCAGGACCCAGACA-3’. The mRNA expression of *Ren1* was normalized to *Rps14* expression. For *Rps14*, the forward primer was 5’-CAGGACCAAGACCCCTGGA-3’, and the reverse was 5’-ATCTTCATCCCAGAGCGAGC-3’. Changes in expression were determined by the ∆∆Ct method and were reported as relative expression compared to control cells.

#### DNA extraction from cells and confirmation of *Nfix* deletion

We extracted DNA from the Trizol fractions remaining after RNA isolation by precipitation with 100% ethanol. The DNA pellet was washed with 0.1 mol/L sodium citrate/10% ethanol followed by 75% ethanol. DNA pellets were dissolved in nuclease-free water. To detect the *Nfix* knockout, we performed PCR using primers: forward: 5’-AGGTCCAGTTCTTTGATTGTGA-3’, reverse: 5’-ACACCTGGTTCAACCTGCAG -3’. We cleaned the PCR reactions using the Qiagen MinElute PCR purification kit (Qiagen). Samples were analyzed by electrophoresis in a 1% agarose gel for the presence of wild-type and mutated fragments. PCR products were subjected to Sanger sequencing using primer 5’-CTGGTTCAACCTGCAGGCGC-3’. We determined the efficiency of *Nfix* deletion from the Sanger sequencing results using the ICE tool from Synthego.

### Ethical approval of animal studies

All animals were handled in accordance with the National Institutes of Health guidelines for the care and use of experimental animals, and the study was approved by the Institutional Animal Care and Use Committee of the University of Virginia.

## Supplemental figures

**Fig. S1:**
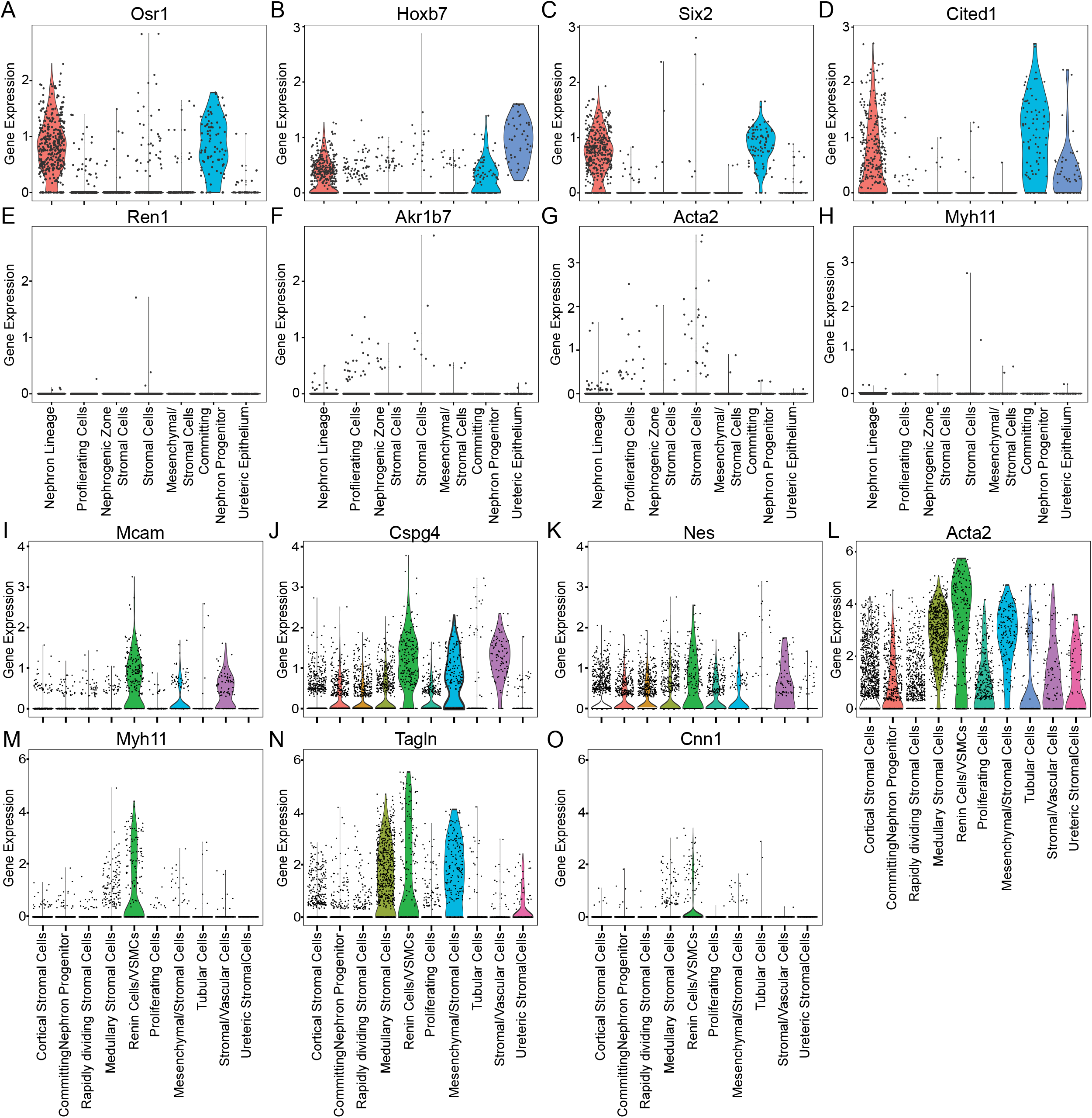
Gene expression levels of different markers among the E12 (A-H) and E18 (I-O) clusters. (A) Osr1 is an intermediate mesoderm marker. Some cells express this marker, meaning they are still at an earlier stage of differentiation. (B) Hoxb7 is an ureteric bud marker. This embryonic compartment gives rise to the collecting ducts in the nephron. Significantly, its highest expression is in the “Ureteric Epithelium” cluster. (C) Six2 and (D) Cited1 are both markers of the cap mesenchyme. This embryonic compartment generates most of the tubular system, podocytes, and Bowman capsule in the nephron. Although we are analyzing FoxD1+ cells, a marker of the loose metanephric mesenchyme, there is some overlap among the embryonic compartments. (E) Renin and (F) Akr1b7 are canonical markers of renin-expressing cells. At E12, they are scarce with low gene expression levels. (G) Smooth Muscle Alpha Actin (Acta2) and (H) Smooth Muscle Myosin Heavy Chain 11 (Myh11) are markers of smooth muscle cells, which remain sparse and with low gene expression at E12. Although no single cluster was labelled pericytes, there are cells with high expression of these markers in two clusters: “Renin Cells/VSMCs” and “Stromal/Vascular Cells.” The pericytes markers are: (I) Melanoma Cell Adhesion Molecule (Mcam), (J) Neuron-glial Antigen 2 (Ng2 or Cspg4), and (K) Nestin (Nes). The VSMCs are spread among all the clusters, as shown by the markers (L) Acta2, (M) Myh11, (N) Tagln, and (O) Cnn1. Despite the widespread distribution of the VSMCs markers, the highest expression is located in the “Renin Cells/VSMCs” cluster. VSMCs: Vascular Smooth Muscle Cells; Cspg4: Chondroitin Sulfate proteoglycan 4.

**Fig. S2:**
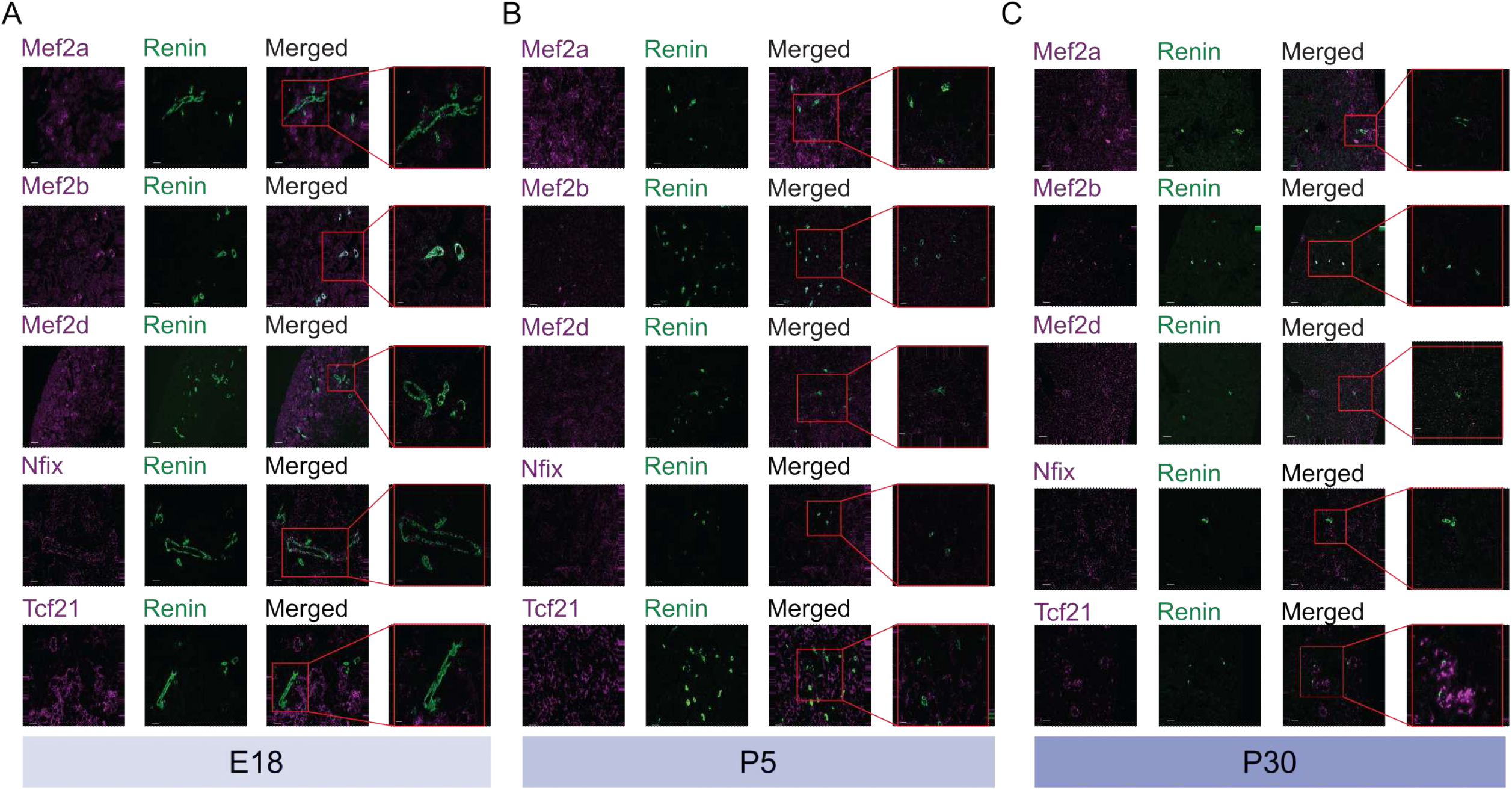
RNAscope of Mef2a, Mef2b, Mef2d, Nfix, and Tcf21 in kidney sections across development. (A) E18. (B) P5. (C) P30. Scale bars: 50µm

**Fig. S3:**
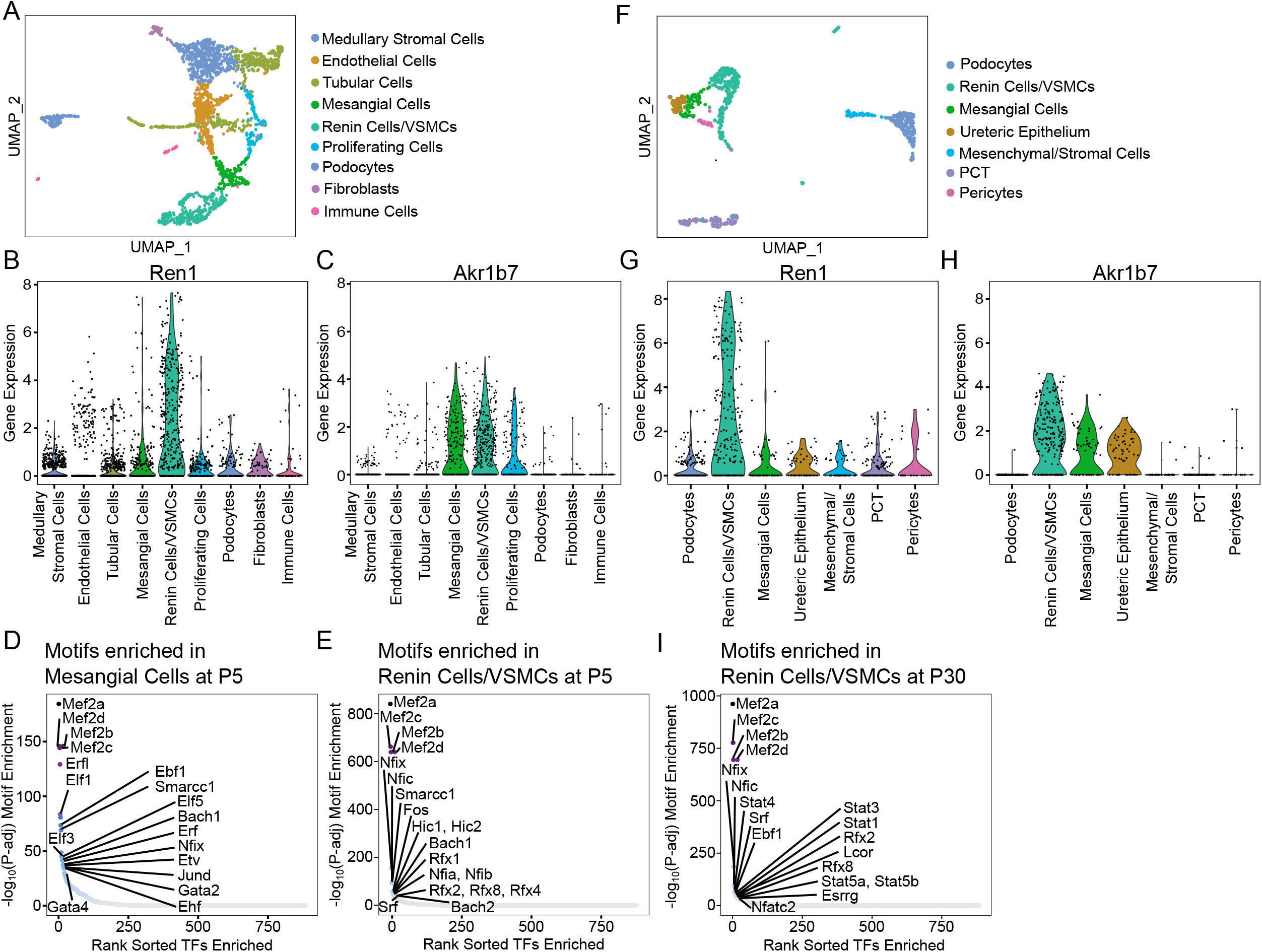
Overview of single time-point analysis at P5 (A-E) and P30 (F-I). (A) UMAP depicting the FoxD1 derivative cells at P5. Violin plots showing Ren1(B) and Akr1b7(C) gene expression levels among the clusters. The “Renin Cells/VSMCs” cluster harbors the highest gene expression level of both markers. The top enriched TF motifs in the (D) “Mesangial Cells” and (E) “Renin Cells/VSMCs” clusters. The MEF2 family of TFs are the highest enriched motifs in both clusters. (F) UMAP depicting the FoxD1 derivative cells at P30. Violin plots showing (G) Ren1 and (H) Akr1b7 gene expression levels among the clusters. The “Renin Cells/VSMCs” cluster shows the highest expression level of both markers. (I) The top enriched motifs in the “Renin Cells/VSMCs”, again highlighting the MEF2 family as the highest enriched motifs.

**Fig. S4:**
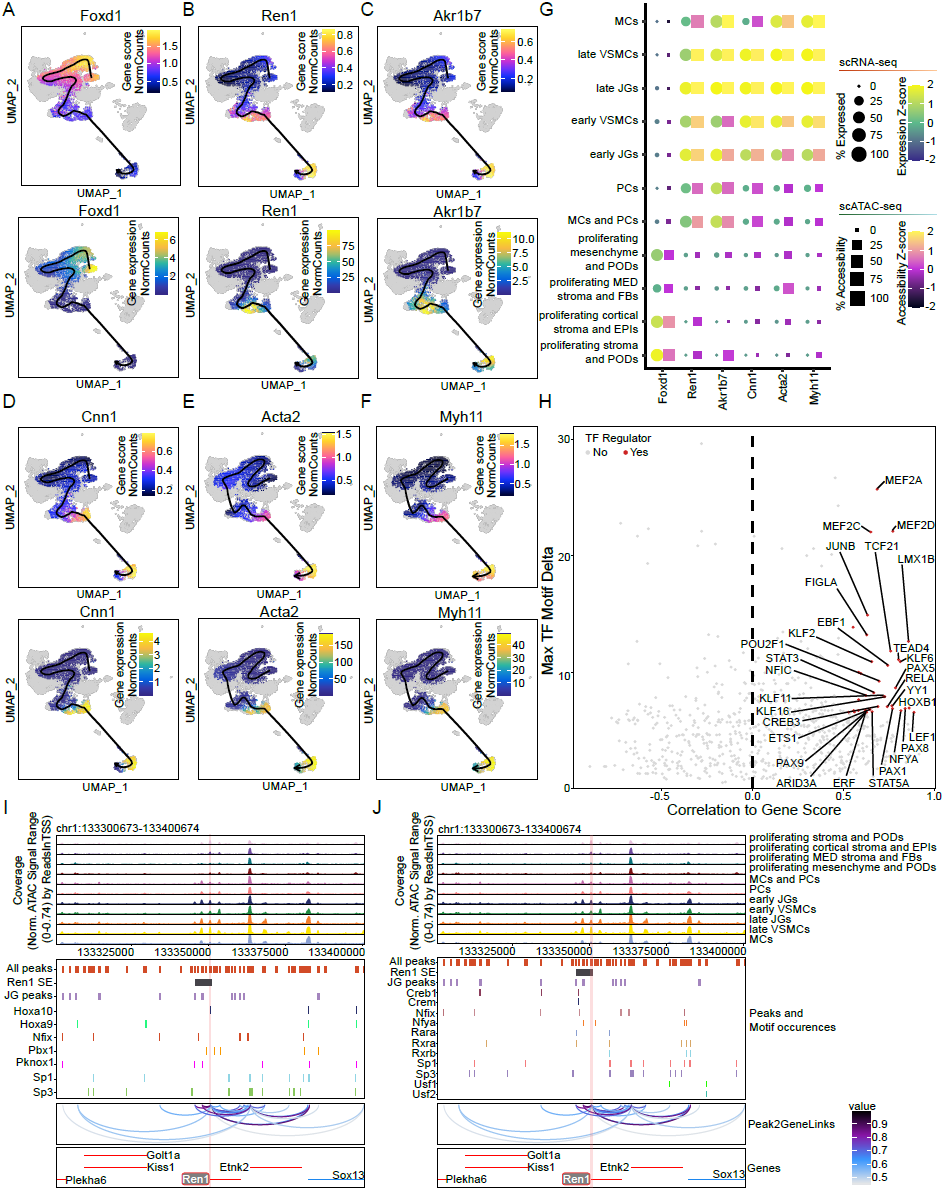
JG cell differentiation trajectory markers and drivers. UMAP visualization of FoxD1 (A), Ren1 (B), Akr1b7 (C), Cnn1 (D), Acta2 (E), and Myh11 (F) gene activity scores (top) and gene expression values (bottom). (G) Dot plot of gene activity scores and gene expression values for each trajectory marker gene. (H) TFs whose gene expression and motif enrichment are positively correlated indicate putative drivers of differentiation. Browser tracks identify preferentially enriched open chromatin in the JG cell cluster at Ren1. Reported promoter binding motif occurrences of enriched TFs identify putative binding sites in open and co-accessible peaks. Pink fill box highlights Ren1 core promoter. (J) Browser tracks identify preferentially enriched open chromatin in the JG cell cluster at Ren1. Reported enhancer binding motif occurrences of enriched TFs identify putative binding sites in open and co-accessible peaks. Pink fill box highlights Ren1 core promoter.

**Fig. S5:**
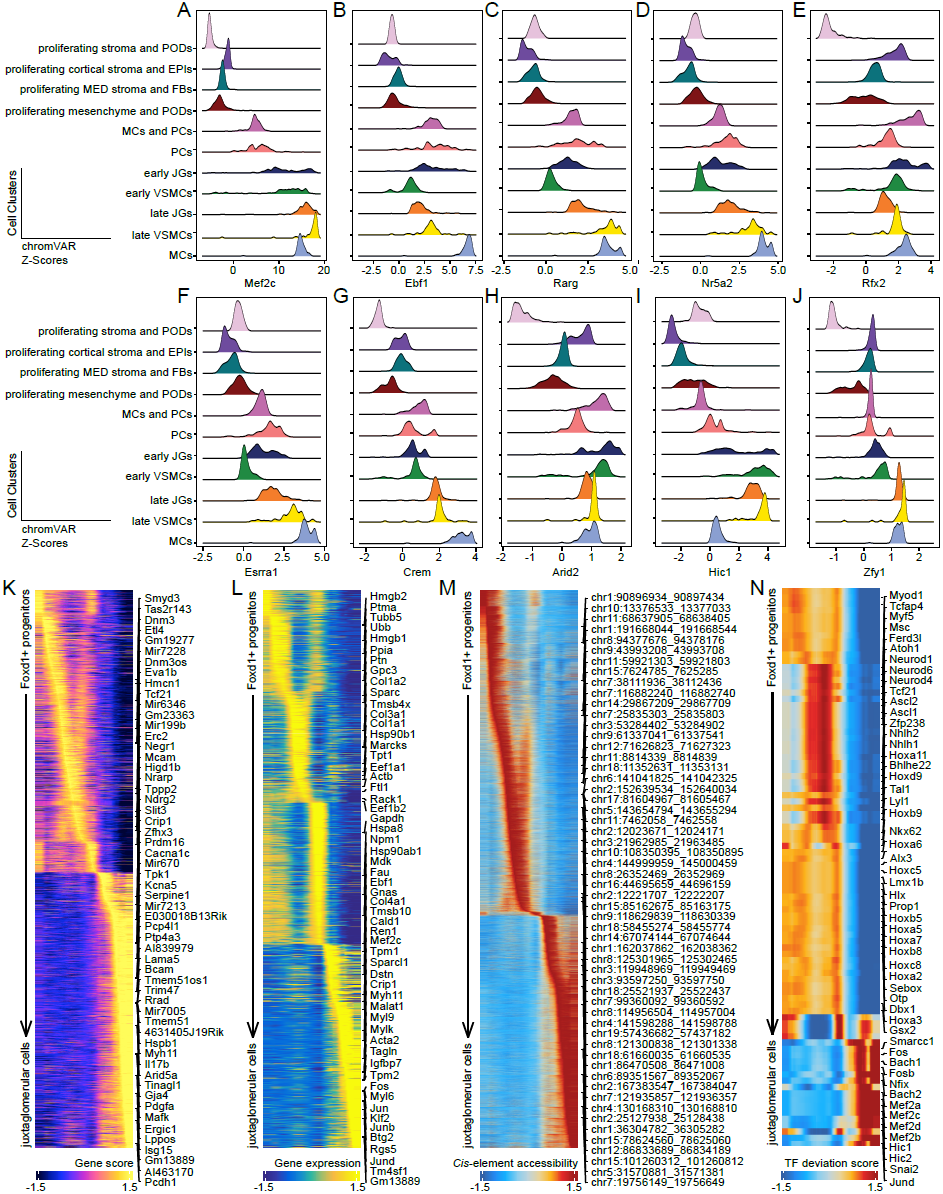
Epigenetic features that drive JG cell differentiation. (A) Mef2c (and Mef2a, Mef2b, and Mef2d (not shown)) are enriched in JG cells and VSMCs. (B) Ebf1 is enriched middle to late and particularly in MCs. (C) Rarg is enriched during the appearance of early JGs before showing overall enrichment late. (D) Nr5a2 is enriched late, particularly in MCs. (E) Rfx2 is positively enriched throughout development but specifically in the early JGs. (F) Esrra1 and (G) Crem become increasingly enriched throughout differentiation. (H) Arid2 is specifically enriched in the early JGs. (I) Hic1 is initially enriched in early JGs and remains enriched in late JGs and VSMCs. (J) Zfy1 becomes enriched as differentiation progresses. Heatmaps of (K) gene score activity, (L) gene expression, (M) accessible chromatin, and (N) enriched TF motifs along the JG trajectory. (EPIs: epithelial cells; FBs: fibroblasts; JG: juxtaglomerular; MED: medullary; MCs: mesangial cells; PCs: pericytes; PODS: podocytes; VSMCs: vascular smooth muscle cells)

**Fig. S6:**
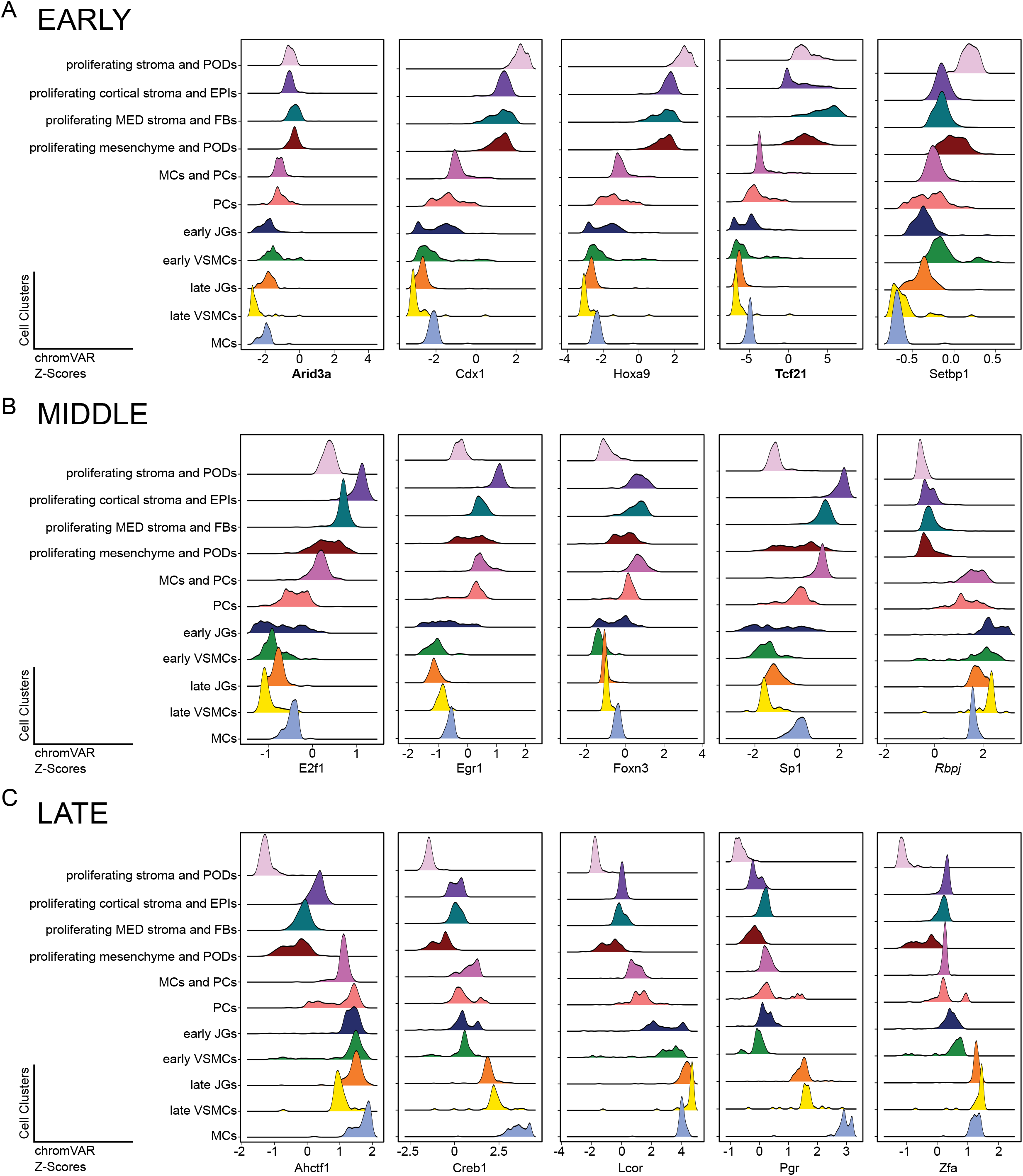
Transcription factors enriched along the JG differentiation trajectory act as early to middle to late acting factors. TFs that are enriched in clusters along the developmental trajectory of JG cells early (A), middle (B), or late (C). **Bold** TFs are positive TF regulators Italicized TFs are known regulators of renin expression. (EPIs: epithelial cells; FBs: fibroblasts; JG: juxtaglomerular; MED: medullary; MCs: mesangial cells; PCs: pericytes; PODS: podocytes; VSMCs: vascular smooth muscle cells)

**Fig. S7:**
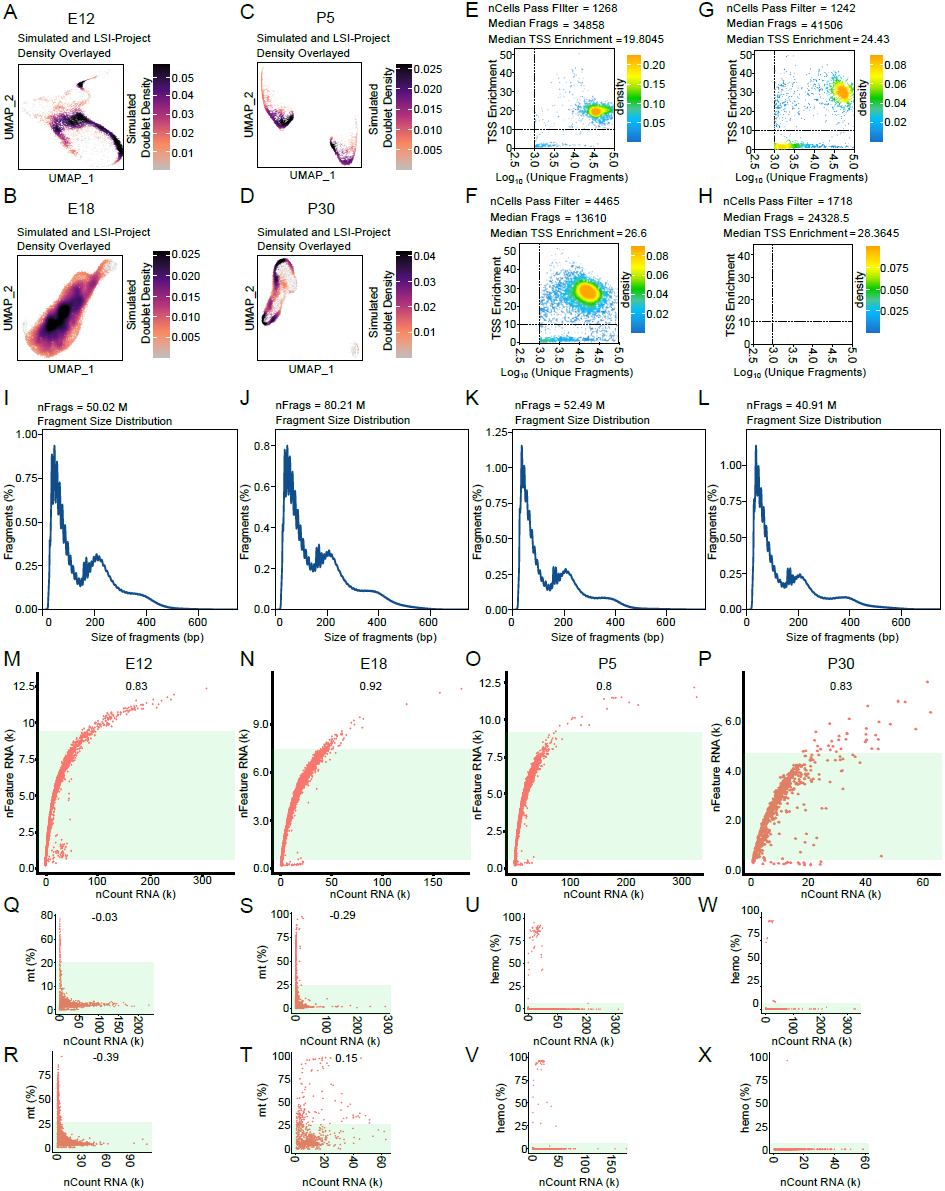
scATAC-seq (A-L) and scRNA-seq (M-X) quality control. UMAP visualiztion of putative doublets in developmental time points E12 (A), E18 (B), P5 (C), and P30 (D). Distributions of TSS enrichment by the log10 number of unique fragments for E12 (E), E18 (F), P5 (G), and P30 (H). Dashed lines represent cut off values below which cells are removed. Fragment distribution plots for developmental time points E12 (I), E18 (J), P5 (K), and P30 (L).Distribution of the number of RNA features against the total RNA count in developmental time points E12 (M), E18 (N), P5 (O), and P30 (P). (e-h) Distribution of the percentage of mitochondria mapped reads against the total RNA count in developmental time points E12 (Q), E18 (R), P5 (S), and P30 (T). Distribution of the percentage of hemoglobin mapped reads against the total RNA count in developmental time points E12 (U), E18 (V), P5 (W), and P30 (X). Green fill boxes represent cells passing filters.

## References

1. Reddi, V., Zaglul, A., Pentz, E.S., and Gomez, R.A. (1998). Renin-expressing cells are associated with branching of the developing kidney vasculature. 9, 63–71. 10.1681/asn.v9163.

2. Starke, C., Betz, H., Hickmann, L., Lachmann, P., Neubauer, B., Kopp, J.B., Sequeira-Lopez, M.L.S., Gomez, R.A., Hohenstein, B., Todorov, V.T., et al. (2015). Renin lineage cells repopulate the glomerular mesangium after injury. Journal of the American Society of Nephrology: JASN 26, 48–54. 10.1681/ASN.2014030265.

3. Pippin, J.W., Sparks, M.A., Glenn, S.T., Buitrago, S., Coffman, T.M., Duffield, J.S., Gross, K.W., and Shankland, S.J. (2013). Cells of renin lineage are progenitors of podocytes and parietal epithelial cells in experimental glomerular disease. The American journal of pathology 183, 542–557. 10.1016/j.ajpath.2013.04.024.

4. Taugner, R., and Hackenthal, E. (1989). The juxtaglomerular apparatus. 10.1007/978-3-642-88426-9.

5. Gomez, R.A., Chevalier, R.L., Carey, R.M., and Peach, M.J. (1990). Molecular biology of the renal renin-angiotensin system. Kidney international. Supplement 30, S18–23.

6. Gomez, R.A. (2017). Fate of renin cells during development and disease. Hypertension (Dallas, Tex. : 1979) 69, 387–395. 10.1161/HYPERTEN-SIONAHA.116.08316.

7. Martinez, M.F., Medrano, S., Brown, E.A., Tufan, T., Shang, S., Bertoncello, N., Guessoum, O., Adli, M., Belyea, B.C., Sequeira-Lopez, M.L.S., et al. (2018). Super-enhancers maintain renin-expressing cell identity and memory to preserve multi-system homeostasis. The Journal of clinical investigation 128, 4787–4803. 10.1172/JCI121361.

8. Nakamura, N., Burt, D.W., Paul, M., and Dzau, V.J. (1989). Negative control elements and cAMP responsive sequences in the tissue-specific expression of mouse renin genes. Proceedings of the National Academy of Sciences of the United States of America 86, 56–59. 10.1073/pnas.86.1.56.

9. Horiuchi, M., Nakamura, N., Tang, S.S., Barrett, G., and Dzau, V.J. (1991). Molecular mechanism of tissue-specific regulation of mouse renin gene expression by cAMP. Identification of an inhibitory protein that binds nuclear transcriptional factor. The Journal of biological chemistry 266, 16247–16254.

10. Borensztein, P., Germain, S., Fuchs, S., Philippe, J., Corvol, P., and Pinet, F. (1994). Cis-regulatory elements and trans-acting factors directing basal and cAMP-stimulated human renin gene expression in chorionic cells. Circulation research 74, 764–773. 10.1161/01.res.74.5.764.

11. Pan, L., Black, T.A., Shi, Q., Jones, C.A., Petrovic, N., Loudon, J., Kane, C., Sigmund, C.D., and Gross, K.W. (2001). Critical roles of a cyclic AMP responsive element and an e-box in regulation of mouse renin gene expression. The Journal of biological chemistry 276, 45530–45538. 10.1074/jbc.M103010200.

12. Klar, J., Sandner, P., Müller, M.W.H., and Kurtz, A. (2002). Cyclic AMP stimulates renin gene transcription in juxtaglomerular cells. Pflugers Archiv: European journal of physiology 444, 335–344. 10.1007/s00424-002-0818-9.

13. Todorov, V.T., Völkl, S., Friedrich, J., Kunz-Schughart, L.A., Hehlgans, T., Vermeulen, L., Haegeman, G., Schmitz, M.L., and Kurtz, A. (2005). Role of CREB1 and NFkappab-p65 in the down-regulation of renin gene expression by tumor necrosis factor alpha. The Journal of biological chemistry 280, 24356–24362. 10.1074/jbc.M502968200.

14. Brunskill, E.W., Sequeira-Lopez, M.L.S., Pentz, E.S., Lin, E., Yu, J., Aronow, B.J., Potter, S.S., and Gomez, R.A. (2011). Genes that confer the identity of the renin cell. Journal of the American Society of Nephrology : JASN 22, 2213–2225. 10.1681/ASN.2011040401.

15. Castellanos Rivera, R.M., Monteagudo, M.C., Pentz, E.S., Glenn, S.T., Gross, K.W., Carretero, O., Sequeira-Lopez, M.L.S., and Gomez, R.A. (2011). Transcriptional regulator RBP-j regulates the number and plasticity of renin cells. Physiological genomics 43, 1021–1028. 10.1152/physiolgenomics.00061.2011.

16. Santos, P.C.J.L., Krieger, J.E., and Pereira, A.C. (2012). Renin-angiotensin system, hypertension, and chronic kidney disease: Pharmacogenetic implications. Journal of pharmacological sciences 120, 77–88. 10.1254/jphs.12r03cr.

17. Castellanos-Rivera, R.M., Pentz, E.S., Lin, E., Gross, K.W., Medrano, S., Yu, J., Sequeira-Lopez, M.L.S., and Gomez, R.A. (2015). Recombination signal binding protein for ig-kJ region regulates juxtaglomerular cell phenotype by activating the myo-endocrine program and suppressing ectopic gene expression. Journal of the American Society of Nephrology : JASN 26, 67–80. 10.1681/ASN.2013101045.

18. Lin, E.E., Pentz, E.S., Sequeira-Lopez, M.L.S., and Gomez, R.A. (2015). Aldo-keto reductase 1b7, a novel marker for renin cells, is regulated by cyclic AMP signaling. American journal of physiology. Regulatory, integrative and comparative physiology 309, R576–84. 10.1152/ajpregu.00222.2015.

19. Petrovic, N., Black, T.A., Fabian, J.R., Kane, C., Jones, C.A., Loudon, J.A., Abonia, J.P., Sigmund, C.D., and Gross, K.W. (1996). Role of proximal promoter elements in regulation of renin gene transcription. The Journal of biological chemistry 271, 22499–22505. 10.1074/jbc.271.37.22499.

20. Pan, L., and Gross, K.W. (2005). Transcriptional regulation of renin: An update. Hypertension (Dallas, Tex. : 1979) 45, 3–8. 10.1161/01.HYP.0000149717.55920.45.

21. Glenn, S.T., Jones, C.A., Gross, K.W., and Pan, L. (2013). Control of renin [corrected] gene expression. Pflugers Archiv : European journal of physiology 465, 13–21. 10.1007/s00424-012-1110-2.

22. Gomez, R.A., and Sequeira-Lopez, M.L.S. (2018). Renin cells in homeostasis, regeneration and immune defence mechanisms. Nature reviews. Nephrology 14, 231–245. 10.1038/nrneph.2017.186.

23. Sequeira-Lopez, M.L.S., and Gomez, R.A. (2021). Renin cells, the kidney, and hypertension. Circulation research 128, 887–907. 10.1161/CIRCRE-SAHA.121.318064.

24. Nagalakshmi, V.K., Li, M., Shah, S., Gigliotti, J.C., Klibanov, A.L., Epstein, F.H., Chevalier, R.L., Gomez, R.A., and Sequeira-Lopez, M.L.S. (2018). Changes in cell fate determine the regenerative and functional capacity of the developing kidney before and after release of obstruction. Clinical science (London, England : 1979) 132, 2519–2545. 10.1042/CS20180623.

25. Tarchini, B., Duboule, D., and Kmita, M. (2006). Regulatory constraints in the evolution of the tetrapod limb anterior-posterior polarity. Nature 443, 985–988. 10.1038/nature05247.

26. Ide, S., Finer, G., Maezawa, Y., Onay, T., Souma, T., Scott, R., Ide, K., Akimoto, Y., Li, C., Ye, M., et al. (2018). Transcription factor 21 is required for branching morphogenesis and regulates the gdnf-axis in kidney development. Journal of the American Society of Nephrology : JASN 29, 2795–2808. 10.1681/ASN.2017121278.

27. Wang, F., Flanagan, J., Su, N., Wang, L.-C., Bui, S., Nielson, A., Wu, X., Vo, H.-T., Ma, X.-J., and Luo, Y. (2012). RNAscope: A novel in situ RNA analysis platform for formalin-fixed, paraffin-embedded tissues. The Journal of molecular diagnostics : JMD 14, 22–29.

28. Berg, A.C., Chernavvsky-Sequeira, C., Lindsey, J., Gomez, R.A., and Sequeira-Lopez, M.L.S. (2013). Pericytes synthesize renin. World journal of nephrology 2, 11–16. 10.5527/wjn.v2.i1.11.

29. Gomez, R.A., Pentz, E.S., Jin, X., Cordaillat, M., and Sequeira Lopez, M.L.S. (2009). CBP and p300 are essential for renin cell identity and morphological integrity of the kidney. American journal of physiology. Heart and circulatory physiology 296, H1255–H1262. 10.1152/ajp-heart.01266.2008.

30. Neubauer, B., Machura, K., Chen, M., Weinstein, L.S., Oppermann, M., Sequeira-Lopez, M.L., Gomez, R.A., Schnermann, J., Castrop, H., Kurtz, A., et al. (2009). Development of vascular renin expression in the kidney critically depends on the cyclic AMP pathway. American journal of physiology. Renal physiology 296, F1006–F1012. 10.1152/ajprenal.90448.2008.

31. Angel, P., and Karin, M. (1991). The role of jun, fos and the AP-1 complex in cell-proliferation and transformation. Biochimica et biophysica acta 1072, 129–157. 10.1016/0304-419x(91)90011-9.

32. Martínez-Zamudio, R.I., Roux, P.-F., Freitas, J.A.N.L.F. de, Robinson, L., Doré, G., Sun, B., Belenki, D., Milanovic, M., Herbig, U., Schmitt, C.A., et al. (2020). AP-1 imprints a reversible transcriptional programme of senescent cells. Nature cell biology 22, 842–855. 10.1038/s41556-020-0529-5.

33. Brunskill, E.W., Georgas, K., Rumballe, B., Little, M.H., and Potter, S.S. (2011). Defining the molecular character of the developing and adult kidney podocyte. PloS one 6, e24640. 10.1371/journal.pone.0024640.

34. Wu, H., Villalobos, R.G., Yao, X., Reilly, D., Chen, T., Rankin, M., Myshkin, E., Breyer, M.D., and Humphreys, B.D. (2022). Mapping the single-cell transcriptomic response of murine diabetic kidney disease to therapies. Cell Metabolism. https://doi.org/10.1016/j.cmet.2022.05.010.

35. Uhlén, M., Fagerberg, L., Hallström, B.M., Lindskog, C., Oksvold, P., Mardinoglu, A., Sivertsson, Å., Kampf, C., Sjöstedt, E., Asplund, A., et al. (2015). Proteomics. Tissue-based map of the human proteome. Science (New York, N.Y.) 347, 1260419. 10.1126/science.1260419.

36. Rossi, G., Taglietti, V., and Messina, G. (2017). Targeting nfix to fix muscular dystrophies. Cell stress 2, 17–19. 10.15698/cst2018.01.121.

37. Martynoga, B., Mateo, J.L., Zhou, B., Andersen, J., Achimastou, A., Urbán, N., Berg, D. van den, Georgopoulou, D., Hadjur, S., Wittbrodt, J., et al. (2013). Epigenomic enhancer annotation reveals a key role for NFIX in neural stem cell quiescence. Genes & development 27, 1769–1786. 10.1101/gad.216804.113.

38. Pan, L., Glenn, S.T., Jones, C.A., Gronostajski, R.M., and Gross, K.W. (2003). Regulation of renin enhancer activity by nuclear factor i and Sp1/Sp3. Biochimica et biophysica acta 1625, 280–290. 10.1016/s0167-4781(03)00016-2.

39. Yang, C.-Y., Ramamoorthy, S., Boller, S., Rosenbaum, M., Gil, A.R., Mittler, G., Imai, Y., Kuba, K., and Grosschedl, R. (2016). Interaction of CCR4–NOT with EBF1 regulates genespecific transcription and mRNA stability in b lymphopoiesis. Genes Dev 30, 2310–2324. 10.1101/gad.285452.116.

40. Belyea, B.C., Xu, F., Pentz, E.S., Medrano, S., Li, M., Hu, Y., Turner, S., Legallo, R., Jones, C.A., Tario, J.D., et al. (2014). Identification of renin progenitors in the mouse bone marrow that give rise to b-cell leukaemia. Nature communications 5, 3273. 10.1038/ncomms4273.

41. Braun, D.A., Warejko, J.K., Ashraf, S., Tan, W., Daga, A., Schneider, R., Hermle, T., Jobst-Schwan, T., Widmeier, E., Majmundar, A.J., et al. (2019). Genetic variants in the LAMA5 gene in pediatric nephrotic syndrome. Nephrology, dialysis, transplantation : official publication of the European Dialysis and Transplant Association - European Renal Association 34, 485–493. 10.1093/ndt/gfy028.

42. Savige, J., and Harraka, P. (2021). Pathogenic LAMA5 variants and kidney disease. In Kidney 360, pp. 1876–1879. 10.34067/KID.0007312021.

43. Watanabe, H., Martini, A.G., Brown, E.A., Liang, X., Medrano, S., Goto, S., Narita, I., Arend, L.J., Sequeira-Lopez, M.L.S., and Gomez, R.A. (2021). Inhibition of the renin-angiotensin system causes concentric hypertrophy of renal arterioles in mice and humans. JCI Insight 6. 10.1172/jci.insight.154337.

44. Saris, J.J., Hoen, P.A.C. ‘t Garrelds, I.M., Dekkers, D.H.W., Dunnen, J.T. den, Lamers, J.M.J., and Jan Danser, A.H. (2006). Prorenin induces intracellular signaling in cardiomyocytes independently of angiotensin II. Hypertension (Dallas, Tex. : 1979) 48, 564–571. 10.1161/01.HYP.0000240064.19301.1b.

45. Beierwaltes, W.H. (2010). The role of calcium in the regulation of renin secretion. American journal of physiology. Renal physiology 298, F1–F11. 10.1152/ajprenal.00143.2009.

46. Sequeira Lopez, M.L.S., and Gomez, R.A. (2010). Novel mechanisms for the control of renin synthesis and release. Current hypertension reports 12, 26–32. 10.1007/s11906-009-0080-z.

47. Sanchez-Niño, M.D., Sanz, A.B., Sanchez-Lopez, E., Ruiz-Ortega, M., Benito-Martin, A., Saleem, M.A., Mathieson, P.W., Mezzano, S., Egido, J., and Ortiz, A. (2012). HSP27/HSPB1 as an adaptive podocyte antiapoptotic protein activated by high glucose and angiotensin II. Laboratory investigation; a journal of technical methods and pathology 92, 32–45. 10.1038/labinvest.2011.138.

48. Matsumoto, T., Urushido, M., Ide, H., Ishihara, M., Hamada-Ode, K., Shimamura, Y., Ogata, K., Inoue, K., Taniguchi, Y., Taguchi, T., et al. (2015). Small heat shock protein beta-1 (HSPB1) is upregulated and regulates autophagy and apoptosis of renal tubular cells in acute kidney injury. PloS one 10, e0126229. 10.1371/journal.pone.0126229.

49. Nyati, K.K., Agarwal, R.G., Sharma, P., and Kishimoto, T. (2019). Arid5a regulation and the roles of Arid5a in the inflammatory response and disease. Frontiers in immunology 10, 2790. 10.3389/fimmu.2019.02790.

50. Nyati, K.K., Masuda, K., Zaman, M.M.-U., Dubey, P.K., Millrine, D., Chalise, J.P., Higa, M., Li, S., Standley, D.M., Saito, K., et al. (2017). TLR4-induced NF-kappaB and MAPK signaling regulate the IL-6 mRNA stabilizing protein Arid5a. Nucleic Acids Research 45, 2687–2703. 10.1093/nar/gkx064.

51. Su, J., Liu, X., Xu, C., Lu, X., Wang, F., Fang, H., Lu, A., Qiu, Q., Li, C., and Yang, T. (2017). NF-kappab-dependent upregulation of (pro)renin receptor mediates high-NaCl-induced apoptosis in mouse inner medullary collecting duct cells. American journal of physiology. Cell physiology 313, C612–C620. 10.1152/ajpcell.00068.2017.

52. Simons, M., Bader, M., and Müller, D.N. (2020). The (pro)renin receptor: What’s in a name? Nature Reviews Nephrology 16, 304–304. 10.1038/s41581-020-0274-9.

53. Le Gal, L., Pellegrin, M., Santoro, T., Mazzolai, L., Kurtz, A., Meda, P., Wagner, C., and Haefliger, J.-A. (2019). Connexin37-dependent mechanisms selectively contribute to modulate angiotensin II -mediated hypertension. Journal of the American Heart Association 8, e010823. 10.1161/JAHA.118.010823.

54. Lozić, M., Filipović, N., Jurić, M., Kosović, I., Benzon, B., Š;olić, I., Kelam, N., Racetin, A., Watanabe, K., Katsuyama, Y., et al. (2021). Alteration of Cx37, Cx40, Cx43, Cx45, Panx1, and renin expression patterns in postnatal kidneys of Dab1-/-(yotari) mice. International journal of molecular sciences 22. 10.3390/ijms22031284.

55. Boor, P., Ostendorf, T., and Floege, J. (2014). PDGF and the progression of renal disease. Nephrology Dialysis Transplantation 29, i45–i54. 10.1093/ndt/gft273.

56. Ostendorf, T., Boor, P., Roeyen, C.R.C. van, and Floege, J. (2014). Platelet-derived growth factors (PDGFs) in glomerular and tubulointerstitial fibrosis. Kidney international supplements 4, 65–69. 10.1038/kisup.2014.12.

57. Igarashi, K., Kurosaki, T., and Roychoudhuri, R. (2017). BACH transcription factors in innate and adaptive immunity. Nature reviews. Immunology 17, 437–450. 10.1038/nri.2017.26.

58. Jang, E., Kim, U.K., Jang, K., Song, Y.S., Cha, J.-Y., Yi, H., and Youn, J. (2019). Bach2 deficiency leads autoreactive b cells to produce IgG autoantibodies and induce lupus through a t cell-dependent extrafollicular pathway. Experimental & Molecular Medicine 51, 1–13. 10.1038/s12276-019-0352-x.

59. Kurokawa, H., Motohashi, H., Sueno, S., Kimura, M., Takagawa, H., Kanno, Y., Yamamoto, M., and Tanaka, T. (2009). Structural basis of alternative DNA recognition by maf transcription factors. Molecular and cellular biology 29, 6232–6244. 10.1128/MCB.00708-09.

60. Germain, S., Konoshita, T., Philippe, J., Corvol, P., and Pinet, F. (1996). Transcriptional induction of the human renin gene by cyclic AMP requires cyclic AMP response element-binding protein (CREB) and a factor binding a pituitary-specific trans-acting factor (Pit-1) motif. Biochemical Journal 316, 107–113. 10.1042/bj3160107.

61. Moriguchi, T., Hamada, M., Morito, N., Terunuma, T., Hasegawa, K., Zhang, C., Yokomizo, T., Esaki, R., Kuroda, E., Yoh, K., et al. (2006). MafB is essential for renal development and F4/80 expression in macrophages. Molecular and cellular biology 26, 5715–5727. 10.1128/MCB.00001-06.

62. Sato, Y., Tsukaguchi, H., Morita, H., Higasa, K., Tran, M.T.N., Hamada, M., Usui, T., Morito, N., Horita, S., Hayashi, T., et al. (2018). A mutation in transcription factor MAFB causes focal segmental glomerulosclerosis with duane retraction syndrome. Kidney international 94, 396–407. 10.1016/j.kint.2018.02.025.

63. Wang, N., Yuan, J., Liu, F., Wei, J., Liu, Y., Xue, M., and Dong, R. (2021). NFIB promotes the migration and progression of kidney renal clear cell carcinoma by regulating PINK1 transcription. PeerJ 9, e10848. 10.7717/peerj.10848.

64. Lu, W., Quintero-Rivera, F., Fan, Y., Alkuraya, F.S., Donovan, D.J., Xi, Q., Turbe-Doan, A., Li, Q.-G., Campbell, C.G., Shanske, A.L., et al. (2007). NFIA haploinsufficiency is associated with a CNS malformation syndrome and urinary tract defects. PLoS genetics 3, e80. 10.1371/journal.pgen.0030080.

65. Tamahara, T., Ochiai, K., Muto, A., Kato, Y., Sax, N., Matsumoto, M., Koseki, T., and Igarashi, K. (2017). The mTOR-Bach2 cascade controls cell cycle and class switch recombination during b cell differentiation. Molecular and cellular biology 37. 10.1128/MCB.00418-17.

66. Miura, Y., Morooka, M., Sax, N., Roychoudhuri, R., Itoh-Nakadai, A., Brydun, A., Funayama, R., Nakayama, K., Satomi, S., Matsumoto, M., et al. (2018). Bach2 promotes b cell receptor–induced proliferation of b lymphocytes and represses cyclin-dependent kinase inhibitors. The Journal of Immunology 200, 2882–2893. 10.4049/jimmunol.1601863.

67. Lin, C., Song, W., Bi, X., Zhao, J., Huang, Z., Li, Z., Zhou, J., Cai, J., and Zhao, H. (2014). Recent advances in the ARID family: Focusing on roles in human cancer. OncoTargets and therapy 7, 315–324.

68. Nyati, K.K., and Kishimoto, T. (2021). Recent advances in the role of Arid5a in immune diseases and cancer. Frontiers in immunology 12, 827611.

69. Zhang, J., Hou, S., You, Z., Li, G., Xu, S., Li, X., Zhang, X., Lei, B., and Pang, D. (2021). Expression and prognostic values of ARID family members in breast cancer. Aging 13, 5621–5637.

70. Suzuki, N., Hirano, K., Ogino, H., and Ochi, H. (2019). Arid3a regulates nephric tubule regeneration via evolutionarily conserved regeneration signal-response enhancers. eLife 8.

71. Martini, A.G., and Danser, A.H.J. (2017). Juxtaglomerular cell phenotypic plasticity. High blood pressure & cardiovascular prevention : the official journal of the Italian Society of Hypertension 24, 231–242.

72. Inoshita, S., Terada, Y., Nakashima, O., Kuwahara, M., Sasaki, S., and Marumo, F. (1999). Roles of E2F1 in mesangial cell proliferation in vitro. Kidney International 56, 2085–2095. https://doi.org/10.1046/j.1523-1755.1999.00799.x.

73. Ledru, N. (2022). RENIN: Regulatory network inference with single cell multiomic data (GitHub).

74. Kato, Y., Kravchenko, V.V., Tapping, R.I., Han, J., Ulevitch, R.J., and Lee, J.D. (1997). BMK1/ERK5 regulates serum-induced early gene expression through transcription factor MEF2C. The EMBO journal 16, 7054–7066. 10.1093/em-boj/16.23.7054.

75. Sacilotto, N., Chouliaras, K.M., Nikitenko, L.L., Lu, Y.W., Fritzsche, M., Wallace, M.D., Nornes, S., García-Moreno, F., Payne, S., Bridges, E., et al. (2016). MEF2 transcription factors are key regulators of sprouting angiogenesis. Genes & development 30, 2297–2309. 10.1101/gad.290619.116.

76. Paciorkowski, A.R., Traylor, R.N., Rosenfeld, J.A., Hoover, J.M., Harris, C.J., Winter, S., Lacassie, Y., Bialer, M., Lamb, A.N., Schultz, R.A., et al. (2013). MEF2C haploinsufficiency features consistent hyperkinesis, variable epilepsy, and has a role in dorsal and ventral neuronal developmental pathways. Neurogenetics 14, 99–111. 10.1007/s10048-013-0356-y.

77. Vrečar, I., Innes, J., Jones, E.A., Kingston, H., Reardon, W., Kerr, B., Clayton-Smith, J., and Douzgou, S. (2017). Further clinical delineation of the MEF2C haploinsufficiency syndrome: Report on new cases and literature review of severe neurodevelopmental disorders presenting with seizures, absent speech, and involuntary movements. Journal of pediatric genetics 6, 129–141. 10.1055/s-0037-1601335.

78. Sugiaman-Trapman, D., Vitezic, M., Jouhilahti, E.-M., Mathelier, A., Lauter, G., Misra, S., Daub, C.O., Kere, J., and Swoboda, P. (2018). Characterization of the human RFX transcription factor family by regulatory and target gene analysis. BMC genomics 19, 181. 10.1186/s12864-018-4564-6.

79. Laurençon, A., Dubruille, R., Efimenko, E., Grenier, G., Bissett, R., Cortier, E., Rolland, V., Swoboda, P., and Durand, B. (2007). Identification of novel regulatory factor x (RFX) target genes by comparative genomics in drosophila species. Genome biology 8, R195. 10.1186/gb-2007-8-9-r195.

80. Barbaux, S., Niaudet, P., Gubler, M.C., Grünfeld, J.P., Jaubert, F., Kuttenn, F., Fékété, C.N., Souleyreau-Therville, N., Thibaud, E., Fellous, M., et al. (1997). Donor splice-site mutations in WT1 are responsible for frasier syndrome. Nature genetics 17, 467–470. 10.1038/ng1297-467.

81. Oikawa, T., and Yamada, T. (2003). Molecular biology of the ets family of transcription factors. Gene 303, 11–34. 10.1016/s0378-1119(02)01156-3.

82. Tan, R.J., and Liu, Y. (2012). Matrix metalloproteinases in kidney homeostasis and diseases. American journal of physiology. Renal physiology 302, F1351–F1361. 10.1152/ajprenal.00037.2012.

83. Brenner BM I.I., Dworkin LD (1986). The kidney. In Comprehensive human physiology: From cellular mechanisms to integration, R. F. (eds). Brenner BM, ed. (Saunders), pp. 124–144. 10.1007/978-3-642-60946-674.

84. Mené, P., Simonson, M.S., and Dunn, M.J. (1989). Physiology of the mesangial cell. Physiological reviews 69, 1347–1424. 10.1152/physrev.1989.69.4.1347.

85. Ghayur, M.N., Krepinsky, J.C., and Janssen, L.J. (2008). Contractility of the renal glomerulus and mesangial cells: Lingering doubts and strategies for the future. Medical hypotheses and research : MHR 4, 1–9.

86. Watanabe, H., Belyea, B.C., Paxton, R.L., Li, M., Dzamba, B.J., DeSimone, D.W., Gomez, R.A., and Sequeira-Lopez, M.L.S. (2021). Renin cell baroreceptor, a nuclear mechanotransducer central for homeostasis. Circulation research 129, 262–276. 10.1161/CIRCRESAHA.120.318711.

87. Ying, L., Morris, B.J., and Sigmund, C.D. (1997). Transactivation of the human renin promoter by the cyclic AMP/protein kinase a pathway is mediated by both cAMP-responsive element binding protein-1 (CREB)-dependent and CREB-independent mechanisms in calu-6 cells. The Journal of biological chemistry 272, 2412–2420. 10.1074/jbc.272.4.2412.

88. Wu, W., Folter, S. de, Shen, X., Zhang, W., and Tao, S. (2011). Vertebrate paralogous MEF2 genes: Origin, conservation, and evolution. PloS one 6, e17334. 10.1371/journal.pone.0017334.

89. Sequeira Lopez, M.L.S., and Gomez, R.A. (2011). Development of the renal arterioles. Journal of the American Society of Nephrology : JASN 22, 2156–2165. 10.1681/ASN.2011080818.

90. Mohamed, T., and Sequeira-Lopez, M.L.S. (2019). Development of the renal vasculature. Seminars in cell & developmental biology 91, 132–146. 10.1016/j.semcdb.2018.06.001.

91. Sequeira López, M.L.S., Pentz, E.S., Nomasa, T., Smithies, O., and Gomez, R.A. (2004). Renin cells are precursors for multiple cell types that switch to the renin phenotype when homeostasis is threatened. Developmental Cell 6, 719–728. https://doi.org/10.1016/S1534-5807(04)00134-0.

92. Rüster, C., and Wolf, G. (2011). Angiotensin II as a morphogenic cytokine stimulating renal fibrogenesis. Journal of the American Society of Nephrology : JASN 22, 1189–1199. 10.1681/ASN.2010040384.

93. Ibrahim, H.N., Jackson, S., Connaire, J., Matas, A., Ney, A., Najafian, B., West, A., Lentsch, N., Ericksen, J., Bodner, J., et al. (2013). Angiotensin II blockade in kidney transplant recipients. Journal of the American Society of Nephrology : JASN 24, 320–327. 10.1681/ASN.2012080777.

94. Cao, W., Li, A., Wang, L., Zhou, Z., Su, Z., Bin, W., Wilcox, C.S., and Hou, F.F. (2015). A salt-induced reno-cerebral reflex activates renin-angiotensin systems and promotes CKD progression. Journal of the American Society of Nephrology : JASN 26, 1619–1633. 10.1681/ASN.2014050518.

95. Zhou, L., Li, Y., Hao, S., Zhou, D., Tan, R.J., Nie, J., Hou, F.F., Kahn, M., and Liu, Y. (2015). Multiple genes of the renin-angiotensin system are novel targets of wnt/b-catenin signaling. Journal of the American Society of Nephrology : JASN 26, 107–120. 10.1681/ASN.2014010085.

96. Kamo, T., Akazawa, H., Suzuki, J.-I., and Komuro, I. (2016). Roles of renin-angiotensin system and wnt pathway in aging-related phenotypes. Inflammation and regeneration 36, 12. 10.1186/s41232-016-0018-1.

97. Perry R. L. S., Yang, C., Soora, N., Salma, J., Marback, M., Naghibi, L., Ilyas, H., Chan, J., Gordon J. W., and McDermott J. C. (2009). Direct interaction between myocyte enhancer factor 2 (MEF2) and protein phosphatase 1α represses MEF2-dependent gene expression. Molecular and Cellular Biology 29, 3355–3366. 10.1128/MCB.00227-08.

98. Kato, Y., Kravchenko, V.V., Tapping, R.I., Han, J., Ulevitch, R.J., and Lee, J.D. (1997). BMK1/ERK5 regulates serum-induced early gene expression through transcription factor MEF2C. The EMBO journal 16, 7054–7066. 10.1093/emboj/16.23.7054.

99. Lin, E.E., Sequeira-Lopez, M.L.S., and Gomez, R.A. (2014). RBP-j in FOXD1+ renal stromal progenitors is crucial for the proper development and assembly of the kidney vasculature and glomerular mesangial cells. American journal of physiology. Renal physiology 306, F249–58. 10.1152/ajprenal.00313.2013.

100. Magaletta, M.E., Lobo, M., Kernfeld, E.M., Aliee, H., Huey, J.D., Parsons, T.J., Theis, F.J., and Maehr, R. (2022). Integration of single-cell transcriptomes and chromatin landscapes reveals regulatory programs driving pharyngeal organ development. Nature Communications 13, 457. 10.1038/s41467-022-28067-4.

101. Marand, A.P., Chen, Z., Gallavotti, A., and Schmitz, R.J. (2021). A cis-regulatory atlas in maize at single-cell resolution. Cell 184, 3041–3055.e21. 10.1016/j.cell.2021.04.014.

102. Martin, M. (2011). Cutadapt removes adapter sequences from high-throughput sequencing reads. EMBnet.journal 17, 10–12. 10.14806/ej.17.1.200.

103. Li, H., and Durbin, R. (2009). Fast and accurate short read alignment with burrows-wheeler transform. Bioinformatics (Oxford, England) 25, 1754–1760. 10.1093/bioinformatics/btp324.

104. Li, H., and Durbin, R. (2010). Fast and accurate long-read alignment with burrows-wheeler transform. Bioinformatics (Oxford, England) 26, 589–595.

105. Granja, J.M., Corces, M.R., Pierce, S.E., Bagdatli, S.T., Choudhry, H., Chang, H.Y., and Greenleaf, W.J. (2021). ArchR is a scalable software package for integrative single-cell chromatin accessibility analysis. Nature genetics 53, 403–411. 10.1038/s41588-021-00790-6.

106. Satpathy, A.T., Granja, J.M., Yost, K.E., Qi, Y., Meschi, F., McDermott, G.P., Olsen, B.N., Mumbach, M.R., Pierce, S.E., Corces, M.R., et al. (2019). Massively parallel single-cell chromatin landscapes of human immune cell development and intratumoral t cell exhaustion. Nature biotechnology 37, 925–936. 10.1038/s41587-019-0206-z.

107. Granja, J.M., Klemm, S., McGinnis, L.M., Kathiria, A.S., Mezger, A., Corces, M.R., Parks, B., Gars, E., Liedtke, M., Zheng, G.X.Y., et al. (2019). Single-cell multiomic analysis identifies regulatory programs in mixed-phenotype acute leukemia. Nature biotechnology 37, 1458–1465. 10.1038/s41587-019-0332-7.

108. Korsunsky, I., Millard, N., Fan, J., Slowikowski, K., Zhang, F., Wei, K., Baglaenko, Y., Brenner, M., Loh, P., and Raychaudhuri, S. (2019). Fast, sensitive and accurate integration of single-cell data with harmony. Nature Methods 16, 1289–1296. 10.1038/s41592-019-0619-0.

109. Zhang, Y., Liu, T., Meyer, C.A., Eeckhoute, J., Johnson, D.S., Bernstein, B.E., Nusbaum, C., Myers, R.M., Brown, M., Li, W., et al. (2008). Model-based analysis of ChIP-seq (MACS). Genome biology 9, R137. 10.1186/gb-2008-9-9-r137.

110. Stuart, T., Butler, A., Hoffman, P., Hafemeister, C., Papalexi, E., Mauck, W.M.3rd., Hao, Y., Stoeckius, M., Smibert, P., and Satija, R. (2019). Comprehensive integration of single-cell data. Cell 177, 1888–1902.e21. 10.1016/j.cell.2019.05.031.

111. McCarthy, D.J., Campbell, K.R., Lun, A.T.L., and Wills, Q.F. (2017). Scater: Pre-processing, quality control, normalization and visualization of single-cell RNA-seq data in r. Bioinformatics (Oxford, England) 33, 1179–1186. 10.1093/bioin-formatics/btw777.

112. Ziegenhain, C., Vieth, B., Parekh, S., Reinius, B., Guillaumet-Adkins, A., Smets, M., Leonhardt, H., Heyn, H., Hellmann, I., and Enard, W. (2017). Comparative analysis of single-cell RNA sequencing methods. Molecular cell 65, 631–643.e4. 10.1016/j.molcel.2017.01.023.

113. Mercer, T.R., Neph, S., Dinger, M.E., Crawford, J., Smith, M.A., Shearwood, A.-M.J., Haugen, E., Bracken, C.P., Rackham, O., Stamatoyannopoulos, J.A., et al. (2011). The human mitochondrial transcriptome. Cell 146, 645–658. 10.1016/j.cell.2011.06.051.

114. AlJanahi, A.A., Danielsen, M., and Dunbar, C.E. (2018). An introduction to the analysis of singlecell RNA-sequencing data. Molecular therapy. Methods & clinical development 10, 189–196. 10.1016/j.omtm.2018.07.003.

115. Hafemeister, C., and Satija, R. (2019). Normalization and variance stabilization of singlecell RNA-seq data using regularized negative binomial regression. Genome biology 20, 296. 10.1186/s13059-019-1874-1.

116. Patil, A., and Patil, A. (2022). CellKb immune: A manually curated database of mammalian hematopoietic marker gene sets for rapid cell type identification. bioRxiv. 10.1101/2020.12.01.389890.

117. Cao, J., Cusanovich, D.A., Ramani, V., Aghamirzaie, D., Pliner, H.A., Hill, A.J., Daza, R.M., McFaline-Figueroa, J.L., Packer, J.S., Christiansen, L., et al. (2018). Joint profiling of chromatin accessibility and gene expression in thousands of single cells. Science 361, 1380–1385. 10.1126/science.aau0730.

118. Park, J., Shrestha, R., Qiu, C., Kondo, A., Huang, S., Werth, M., Li, M., Barasch, J., and Suszták, K. (2018). Single-cell transcriptomics of the mouse kidney reveals potential cellular targets of kidney disease. Science (New York, N.Y.) 360, 758–763. 10.1126/science.aar2131.

119. Combes, A.N., Phipson, B., Lawlor, K.T., Dorison, A., Patrick, R., Zappia, L., Harvey, R.P., Oshlack, A., and Little, M.H. (2019). Single cell analysis of the developing mouse kidney provides deeper insight into marker gene expression and ligand-receptor crosstalk. Development (Cambridge, England) 146. 10.1242/dev.178673.

120. Dijk, D. van, Sharma, R., Nainys, J., Yim, K., Kathail, P., Carr, A.J., Burdziak, C., Moon, K.R., Chaffer, C.L., Pattabiraman, D., et al. (2018). Recovering gene interactions from single-cell data using data diffusion. Cell 174, 716–729.e27. 10.1016/j.cell.2018.05.061.

121. Weirauch, M.T., Yang, A., Albu, M., Cote, A.G., Montenegro-Montero, A., Drewe, P., Najafabadi, H.S., Lambert, S.A., Mann, I., Cook, K., et al. (2014). Determination and inference of eukaryotic transcription factor sequence specificity. Cell 158, 1431–1443. 10.1016/j.cell.2014.08.009.

122. Schep, A.N., Wu, B., Buenrostro, J.D., and Greenleaf, W.J. (2017). chromVAR: Inferring transcription-factor-associated accessibility from single-cell epigenomic data. Nature methods 14, 975–978. 10.1038/nmeth.4401.

123. Grant, C.E., Bailey, T.L., and Noble, W.S. (2011). FIMO: Scanning for occurrences of a given motif. Bioinformatics (Oxford, England) 27, 1017–1018. 10.1093/bioinformatics/btr064.

124. Lawrence, W.A.P., Michael and Huber (2013). Software for computing and annotating genomic ranges. PLOS Computational Biology 9, 1–10. 10.1371/journal.pcbi.1003118.

125. Sigmund, C.D., Okuyama, K., Ingelfinger, J., Jones, C.A., Mullins, J.J., Kane, C., Kim, U., Wu, C.Z., Kenny, L., and Rustum, Y. (1990). Isolation and characterization of renin-expressing cell lines from transgenic mice containing a renin-promoter viral oncogene fusionconstruct. The Journal of biological chemistry 265, 19916–19922.

126. Ma, Z., Lytle, N.K., Ramos, C., Naeem, R.F., and Wahl, G.M. (2022). Single-cell transcriptomic and epigenetic analyses of mouse mammary development starting with the embryo. In Mammary stem cells: Methods and protocols, M. dM. Vivanco, ed. (Springer US), pp. 49–82. 10.1007/978-1-0716-2193-63.

127. Dobin, A., Davis, C.A., Schlesinger, F., Drenkow, J., Zaleski, C., Jha, S., Batut, P., Chaisson, M., and Gingeras, T.R. (2013). STAR: Ultrafast universal RNA-seq aligner. Bioinformatics (Oxford, England) 29, 15–21. 10.1093/bioinformat-ics/bts635.

128. Osorio, D., and Cai, J.J. (2021). Systematic determination of the mitochondrial proportion in human and mice tissues for single-cell RNA-sequencing data quality control. Bioinformatics (Oxford, England) 37, 963–967. 10.1093/bioin-formatics/btaa751.

129. Macosko, E.Z., Basu, A., Satija, R., Nemesh, J., Shekhar, K., Goldman, M., Tirosh, I., Bialas, A.R., Kamitaki, N., Martersteck, E.M., et al. (2015). Highly parallel genome-wide expression profiling of individual cells using nanoliter droplets. 161, 1202–1214. 10.1016/j.cell.2015.05.002.

